# Rad53 controls DNA unwinding after helicase-polymerase uncoupling at DNA replication forks

**DOI:** 10.1101/811141

**Authors:** Sujan Devbhandari, Dirk Remus

**Author notes:** Corresponding author: Molecular Biology Program, Memorial Sloan-Kettering Cancer Center, 1275 York Avenue, New York, NY 10065, USA.

## Abstract

The coordination of DNA unwinding and synthesis at replication forks promotes efficient and faithful replication of chromosomal DNA. Using the reconstituted budding yeast DNA replication system, we demonstrate that Pol ε variants harboring catalytic point mutations in the Pol2 polymerase domain, contrary to Pol2 polymerase domain deletions, inhibit DNA synthesis at replication forks by displacing Pol δ from PCNA/primer-template junctions, causing excessive DNA unwinding by the replicative DNA helicase, CMG, uncoupled from DNA synthesis. Mutations that suppress the inhibition of Pol δ by Pol ε restore viability in Pol2 polymerase point mutant cells. We also observe uninterrupted DNA unwinding at replication forks upon dNTP depletion or chemical inhibition of DNA polymerases, demonstrating that leading strand synthesis is not tightly coupled to DNA unwinding by CMG. Importantly, the Rad53 kinase controls excessive DNA unwinding at replication forks by limiting CMG helicase activity, suggesting a mechanism for fork-stabilization by the replication checkpoint.

## INTRODUCTION

The maintenance of genomes across generations depends on the rapid, complete, and accurate replication of the chromosomal DNA. In eukaryotes, three essential DNA polymerases, Pol α, Pol δ, and Pol ε, are required for normal chromosome replication ^1–3^. The conservation of this multi-polymerase DNA replication strategy in eukaryotes points to a functional specialization of replicative DNA polymerases. Indeed, a large body of evidence has uncovered strand-specific roles for these polymerases, indicating that Pol α and Pol δ act preferentially on the lagging strand, while Pol ε synthesizes the bulk of the leading strand ^2,3^. Reconstitution studies in budding yeast have, moreover, elucidated biochemical and structural mechanisms that target the polymerases to their respective strands ^4^.

Pol2, the catalytic subunit of Pol ε, contains two exonuclease-polymerase domains ^5^. The N-terminal domain is catalytically active, while the C-terminal domain is catalytically inactive ^6,7^. Point mutations in the polymerase active site of Pol2 are lethal, but surprisingly deletion of the catalytic domain is not lethal, suggesting that Pol ε performs an essential non-catalytic function in the cell ^8,9^. One such function involves the assembly of the replicative DNA helicase, CMG (Cdc45-MCM-GINS), at replication origins ^1^. In *pol2-16* cells lacking the Pol2 catalytic domain, leading strand synthesis is largely carried out by Pol δ, but S phase is severely defective in these cells ^10,11^. In the absence of Pol ε polymerase activity, the rate and processivity of leading strand synthesis are compromised *in vitro*, consistent with DNA synthesis by Pol ε being important for normal DNA replication ^12,13^.

Maximal fork rates likely depend on the ability of Pol ε, unlike Pol δ, to physically associate with CMG ^6,14^. Helicase-polymerase coupling significantly enhances fork rates in prokaryotic systems ^15–18^, and recent observations suggest similar cooperativity between CMG and Pol ε at eukaryotic replication forks ^12,13, 19–21^. However, helicase-polymerase coupling is challenged in conditions that inhibit polymerase, but not helicase, progression. In eukaryotes, helicase-polymerase uncoupling has been observed in the presence of leading strand lesions and DNA-protein crosslinks, and in conditions that inhibit DNA polymerase catalytic activity ^20–25^. Single-stranded DNA (ssDNA) generated in the wake of uncoupled CMG plays an important physiological role by providing a platform for the activation of the apical checkpoint kinase, Mec1-Ddc2/ATR-ATRIP (budding yeast/vertebrates) ^26,27^. Downstream activation of the effector kinase, Rad53/CHK1, elicits a host of responses aimed at preserving genome integrity during replication stress, including cell cycle arrest, increased dNTP levels, nuclease inhibition, and inhibition of origin firing ^28–30^.

The critical function of the checkpoint during genotoxic stress is to protect stalled forks from irreversible breakdown ^31^. The mechanism of fork stabilization is obscure and may involve distinct pathways depending on the stress condition, but it does not appear to require stabilization of the replisome itself ^32–35^. However, checkpoint activation restricts replisome progression and minimizes the generation of ssDNA at stalled forks^24,25,32,36^. How replisome progression may be controlled by the checkpoint is not known.

Here we report that Pol ε polymerase point mutants, unlike polymerase domain deletions, inhibit DNA synthesis by Pol δ at replication forks *in vitro* by displacing Pol δ from PCNA-loaded nascent 3’ ends. Mutations that impair the ability of Pol ε to compete Pol δ suppress DNA replication defects *in vitro* and restore viability of Pol2 polymerase point mutant cells. Unexpectedly, inhibition of DNA replication by mutant Pol ε induces excessive DNA unwinding by the CMG helicase. Uninterrupted progression of CMG after stalling of DNA synthesis is a general response of the budding yeast replisome to polymerase inhibition, indicating that DNA unwinding and DNA synthesis are not tightly coupled at eukaryotic forks. Importantly, Rad53 can constrain replisome progression by directly inhibiting the CMG helicase, suggesting a mechanism for fork stabilization by the replication checkpoint.

## RESULTS

### Regulated replication of nucleosome-free plasmid templates *in vitro*

Pol2 polymerase point mutations, unlike Pol2 catalytic domain deletions, are lethal ^8^. Yet, fork rates in the presence of a catalytic mutant of Pol ε (Pol ε^D640A^) or Pol ε lacking the catalytic domain (Pol ε^Δcat^) appeared to be similar *in vitro* ^13,37^. Therefore, to investigate potential differences between these Pol ε variants, we decided to characterize them side-by-side using the reconstituted origin-dependent budding yeast DNA replication system ^13,37^. We had previously demonstrated that nucleosomes limit Okazaki fragment length *in vitro* by restricting strand-displacement synthesis by Pol δ ^37^. However, using higher salt and lower Pol δ concentrations we observe Okazaki fragments with a physiological length distribution also on naked DNA templates (**Figure S1**).

The modified conditions support efficient maturation of nascent strands by Cdc9 and Fen1 and normal replication of plasmid DNA templates (**Figures 1A+B**). In the presence of Top2, the two predominant ligation products were full-length or close to full-length linears (ssL), and covalently closed circles (ccc) (**Figures 1A**). The latter are an expected product of termination, while the former may result from fork stalling upon convergence ^38^. Linear full-length or close to full-length daughter strands were also generated in the presence of Top1. As expected, due to the inability of Top1 to decatenate double-stranded DNA molecules, circular daughter strands remain topologically linked (cat). The ratio of circular to linear daughter strands was reduced in the presence of Top1 compared to Top2, indicating that Top2 has a greater proficiency in promoting replication termination. On average, we observe a termination efficiency of ∼30 % in the presence of Top1 and ∼45 % in the presence of Top2 within 45 minutes after origin firing on 4.8 kbp plasmid templates (**Figure S2A**). In the absence of topoisomerase, replication forks stalled ∼ 500-700 bp from the origin due to the accumulation of positive supercoils, resulting in the appearance of an early replication intermediate (ERI) (**Figures 1B** and S2C). In the presence of Top1, leading strands were fully extended to approximately half-unit length (**Figure S2B**), and the predominant products are a late replication intermediate (LRI) and catenated plasmid dimers, the latter of which appear as a ladder of bands migrating between the nicked monomer and the LRI (**Figure 1B**). Pulse-chase analysis demonstrates that the ERI is a precursor of the LRI (**Figure S2C**). The two most prominent replication products in the presence of Top2 are plasmid monomers and LRIs (**Figure 1B**). At high Top2 concentrations, DNA knots also became apparent (**Figure 1B**). In the presence of both Top1 and Top2, replication products resembled those obtained with Top 2 alone (**Figure S2D**), which is the condition we will use in standard replication reactions below.

**Figure 1:**
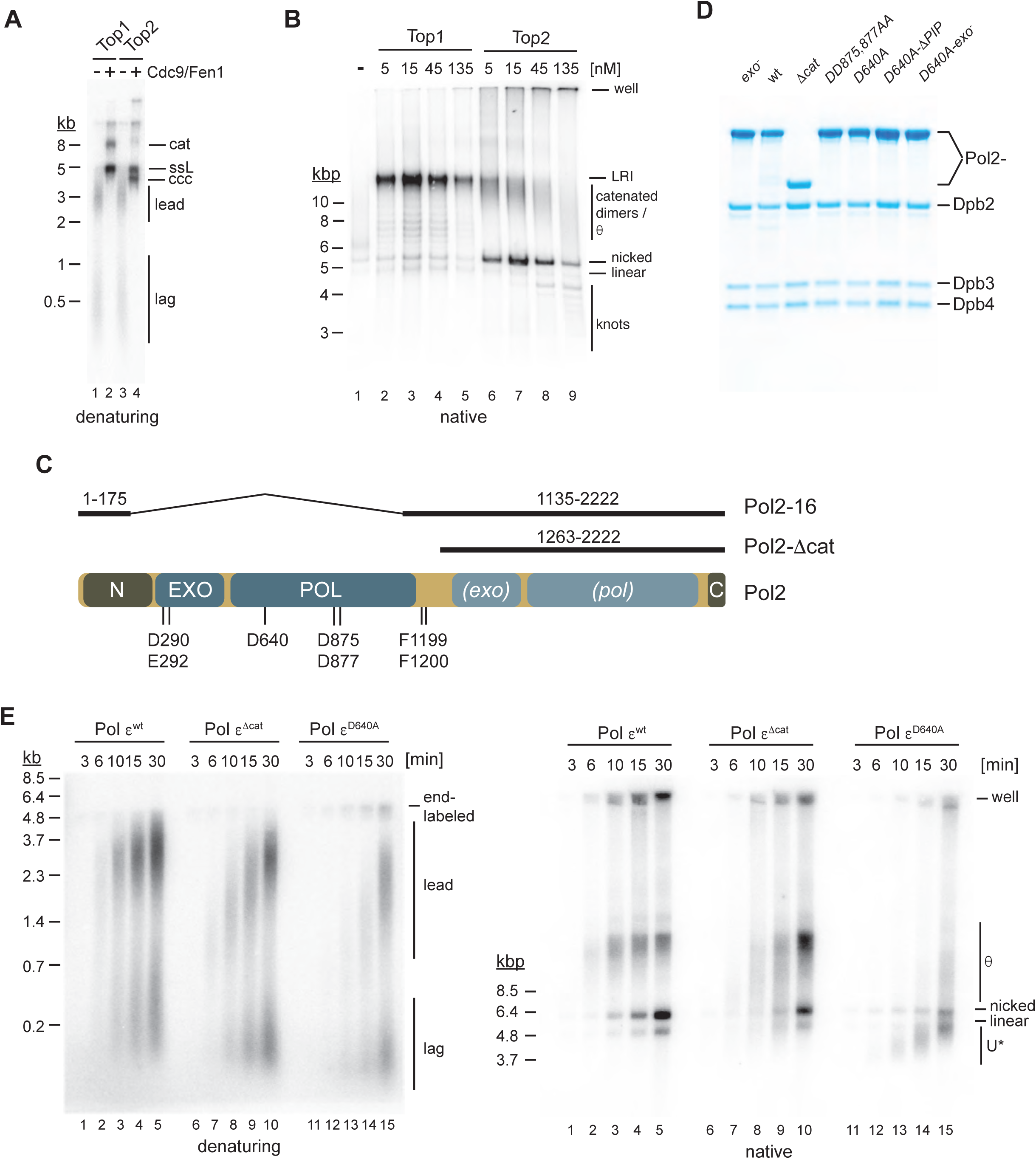
Active site mutations in the Pol ε polymerase domain inhibit DNA synthesis and induce excessive DNA unwinding at replication forks. **(A)** Nascent strand maturation by Cdc9 and Fen1 in standard DNA replication reactions. Template: pARS1. Replication products were analyzed by 0.8 % alkaline agarose gel-electrophoresis and autoradiography. lead: leading strands; lag: lagging strands; ccc: covalently closed circle; ssL: single-stranded linear; cat: catenated dimer. **(B)** Native agarose gel-analysis of standard replication products. Template: pARS1. LRI: late replication intermediate. **(C)** Domain structure of Pol2. Amino acids mutated in this study are indicated at the bottom. Pol2 regions retained in Pol2-16 and Pol2-Δcat are indicated on top. N: N-terminal domain; EXO: exonuclease domain; POL: polymerase domain; *(exo)*: inactive exonuclease domain; *(pol)*: inactive polymerase domain; C: C-terminal domain. **(D)** Purified Pol ε variants. **(E)** Time course analysis of standard DNA replication reactions performed with indicated Pol ε variants (15 nM); template: pARS1. Replication products were analyzed by 0.8 % alkaline (left) or native (right) agarose gel-electrophoresis and autoradiography. U*: partially replicated and unwound DNA.

### Pol ε polymerase point mutants cause extensive DNA unwinding at replication forks

Our attempts to purify Pol ε containing *pol2-16*, which lacks the Pol2 catalytic domain, were largely unsuccessful. We, therefore, purified Pol ε^Δcat^ to study the deletion of the Pol2 catalytic domain ^13^. We also purified Pol ε complexes harboring alanine substitutions at Pol2 residues D640 or DD875,877 to test the effect of point mutations in the Pol2 polymerase domain ^8,37^. Consistent with previous reports, the extent of DNA synthesis in the presence of Pol ε^Δcat^ and Pol ε^D640A^ was reduced relative to Pol ε^wt^ (**Figures 1E and S3**) ^13,37^. As RFC inhibits primer extension by Pol α in the reconstituted system, the bulk of DNA synthesis in the presence of Pol ε^Δcat^ and Pol ε^D640A^ is carried out by Pol δ ^37^. The slow fork progression is consistent with Pol δ replicating DNA in a slow and distributive manner physically uncoupled from the CMG ^12,14^. However, the extent of DNA synthesis was significantly lower in the presence of Pol ε^D640A^ or Pol ε^DD875,877AA^ relative to Pol ε^Δcat^ (**Figures 1E, S3, and S4**), indicating that the presence of an inactive Pol2 polymerase domain further inhibits DNA replication.

Analysis of the native structure of the replication products revealed striking differences. As expected, fully replicated nicked plasmid monomers and LRIs were predominant in the presence of Pol ε^wt^ (**Figure 1E**). The fraction of θ intermediates (see **Figure S5** for definition of θ) was increased in the presence of Pol ε^Δcat^, as expected from the lower rate of fork progression. Unexpectedly, however, products generated in the presence of Pol ε^D640A^ or Pol ε^DD875,877AA^ exhibited a greatly increased gel mobility that was maximal early in the reaction and decreased as DNA synthesis progressed (**Figures 1E** and **S4**). Fast migrating plasmid forms are a product of DNA unwinding, which causes negative template supercoiling after deproteinization ^39^ (**Figure S5**, also see **Figure S9**). We refer to these partially replicated and unwound DNA forms as U* to differentiate them from fully unwound U form DNA described previously in other systems in the absence of DNA synthesis ^39–41^. In conclusion, Pol ε complexes harboring point mutations in the polymerase active site inhibit Pol δ, causing excessive DNA unwinding by CMG at replication forks.

### CMG uncoupling from DNA synthesis is caused by free mutant Pol ε

Pol ε physically associates with CMG at replication forks during leading strand synthesis at the fork ^6,12,14, 42–45^. Our observation, however, that leading strand synthesis by Pol δ in the presence of Pol ε polymerase point mutants lagged far behind the advancing CMG suggested that Pol δ was inhibited at a distance behind the fork by free Pol ε unbound from CMG. Consistent with this notion, leading strand lengths correlated inversely with the concentration of Pol ε^D640A^. While leading strands reached normal half-unit length in the presence of 7.5 nM Pol ε^D640A^, their average length decreased as the concentration of Pol ε^D640A^ was increased, reaching about half maximal length at 60 nM Pol ε^D640A^ (**Figure 2A**). This effect is explained by changes in the rate of leading strand synthesis by Pol δ: Using pulse-chase analysis, we observe a rate of leading strand synthesis of 0.28 kb/min in the presence of 7.5 nM Pol ε^D640A^, but only 0.04 kb/min in the presence of 60 nM Pol ε^D640A^ (**Figure 2B**). Formation of U* was also dependent on the concentration of Pol ε^D640A^: While normal θ-type replication intermediates predominated in the presence of 7.5 nM and 15 nM Pol ε^D640A^, U* intermediates were the major products at 30 nM and 60 nM Pol ε^D640A^ (**Figure 2A**).

**Figure 2:**
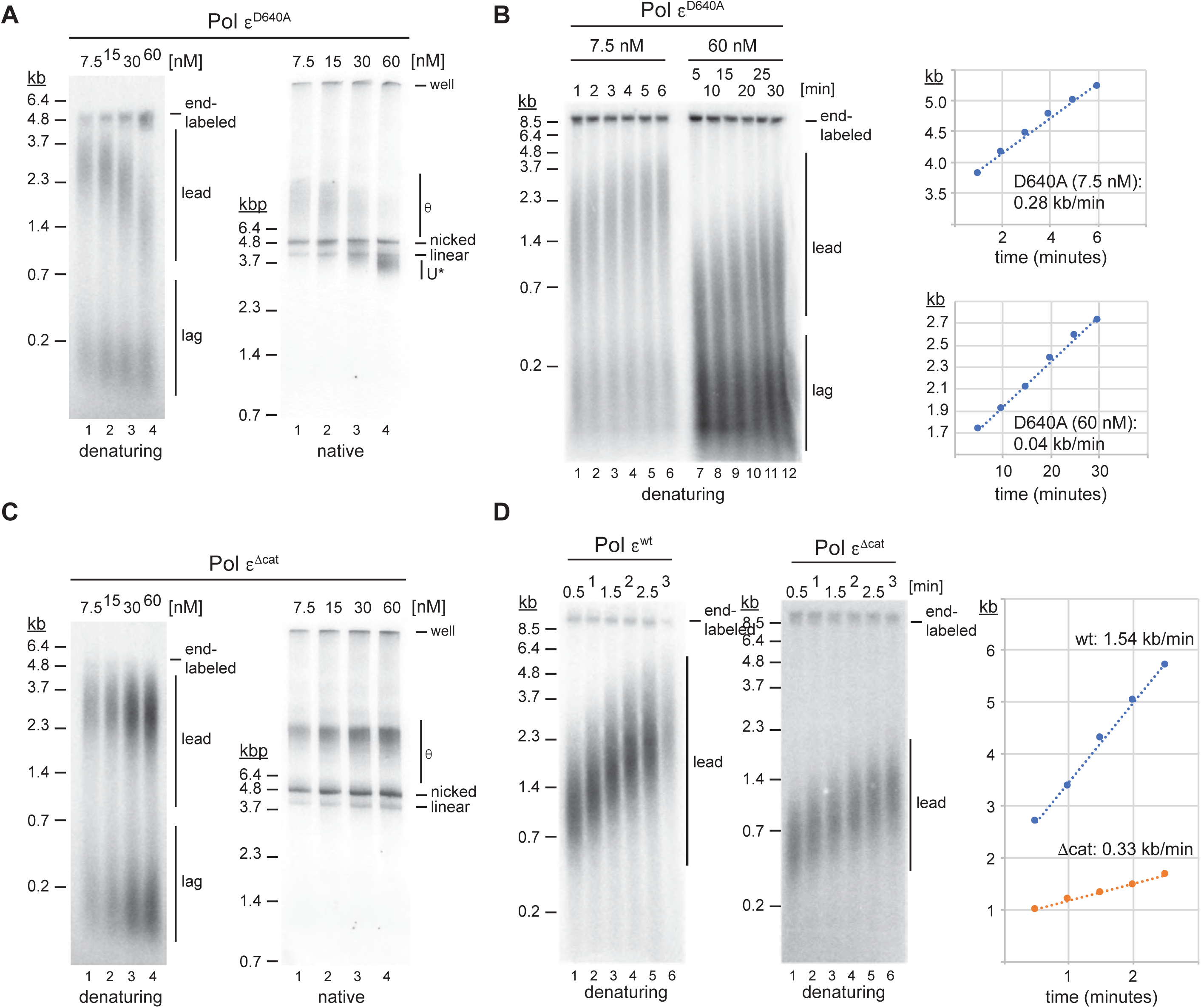
Pol ε polymerase point mutants impede leading strand synthesis in a concentration-dependent manner. **(A)** Titration experiment to determine the effect of Pol ε^D640A^ concentration on fork progression; reactions were stopped 45 minutes after origin activation. Template: pARS1. **(B)** Pulse-chase experiments to determine leading strand synthesis rates in the presence of 7.5 nM or 60 nM Pol ε^D640A^; time indicates minutes after chase with cold dATP; maximum leading strand positions are plotted on the right. Template: pARS305. **(C)** Titration experiment to determine effect of Pol ε^Δcat^ concentration on fork progression. Template: pARS1. **(D)** Pulse-chase experiment to determine rates of leading strand synthesis in the presence of 30 nM Pol ε^wt^ (left) or Pol ε^Δcat^ (middle); time indicates minutes after chase. Template: pARS305. Maximum leading strand positions are plotted on the right.

In contrast, leading strand synthesis rates appeared constant across a range of Pol ε^wt^ and Pol ε^Δcat^ concentrations, and U* replication products were not observed (**Figures 2C and S6**). Consistent with previous reports ^13^, the rate of leading strand synthesis in the presence of Pol ε^Δcat^ was significantly reduced compared to that in the presence of Pol ε^wt^ (0.33 kb/min vs 1.54 kb/min, **Figure 2D**), but very similar to that at low concentrations of Pol ε^D640A^ (0.28 kb/min, **Figure 2B**). The comparable rates of leading strand synthesis and the lack of extensive template unwinding in the presence of either Pol ε^Δcat^ or low concentrations of Pol ε^D640A^ indicate that in both conditions Pol δ extends leading strands in close proximity to CMG.

### Pol ε polymerase point mutants compete with Pol δ for nascent 3’ ends

The competition of Pol δ by mutant Pol ε was unexpected as previous studies had demonstrated that Pol δ dominates over free Pol ε for the binding to PCNA/primer-template junctions ^12,14,46^. To clarify this apparent discrepancy, we tested the ability of mutant and wild-type Pol ε complexes to displace Pol δ from PCNA/primer-template junctions in primer extension assays. Pol δ (4 nM) was pre-incubated with singly-primed RPA-coated M13 ssDNA (1 nM), RFC/PCNA, and three nucleotides to initiate primer extension (**Figure 3A**). Variable concentrations of wild-type or mutant Pol ε were subsequently added along with the remaining fourth nucleotide and DNA synthesis monitored by denaturing gel-analysis (**Figure 3B**). DNA synthesis was allowed to proceed for 3 minutes, which was sufficient time for Pol δ to complete the synthesis of full-length daughter strands, but which yielded only incomplete DNA products by Pol ε under these conditions (**Figure S7**). Competition of Pol δ by wild-type or mutant Pol ε is, therefore, expected to yield shorter than full-length DNA products.

**Figure 3:**
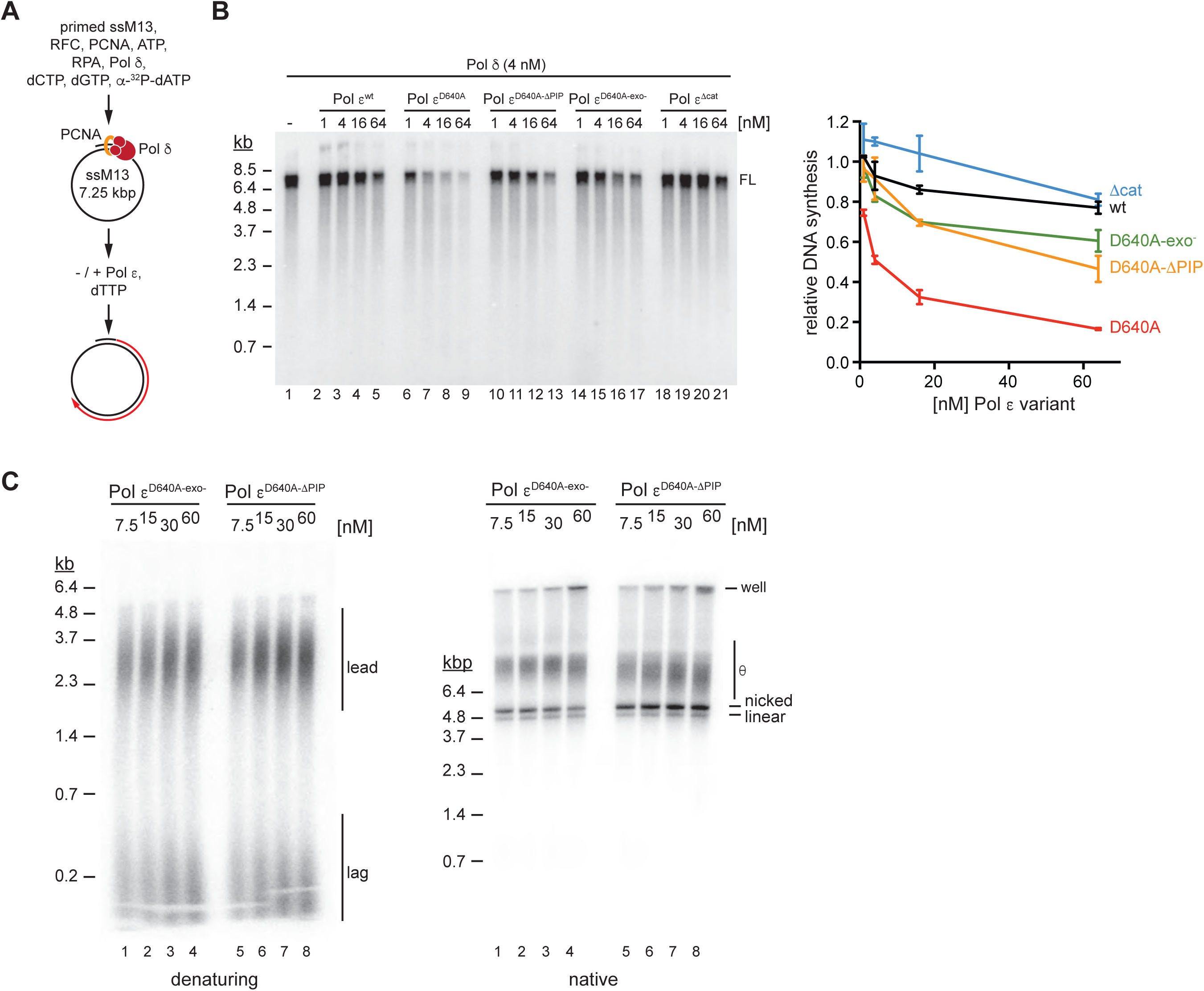
PCNA binding and exonuclease activity of Pol2 are required for inhibition of Pol δ by Pol ε. **(A)** Primer extension reaction scheme to measure Pol δ competition by Pol ε. **(B)** Left: Denaturing agarose gel analysis of primer extension products obtained with PCNA/Pol δ in presence of increasing concentrations of wild-type or mutant Pol ε. FL: full-length (7.25 kb). Right: Average DNA synthesis, normalized to DNA synthesis by Pol δ in absence of Pol ε; error bars indicate standard deviation of two independent experiments. **(C)** Titration experiments to determine the effect of Pol ε^D640A-exo-^ or Pol ε^D640A-ΔPIP^ concentration on fork progression. Reactions were stopped 45 minutes after origin activation. Template: pARS1.

Consistent with a previous report ^12^ we observed that Pol δ was resistant to competition by Pol ε^wt^ over a range of concentrations (**Figure 3B**); even at a 16-fold excess of Pol ε^wt^ over Pol δ DNA synthesis was only reduced by ∼ 20 %. In contrast, a ∼ 50 % reduction in DNA synthesis by Pol δ was observed at equimolar concentrations of Pol ε^D640A^, which increased to a reduction by ∼ 80 % at 16-fold molar excess of Pol ε^D640A^. On the other hand, Pol δ was largely resistant to competition by Pol ε^Δcat^. Thus, Pol ε harboring a point mutation in the polymerase active site, unlike wild-type Pol ε or Pol ε lacking the polymerase domain, exhibits an enhanced capacity to compete Pol δ, providing an explanation for the inhibition of DNA synthesis at replication forks in the presence of Pol ε^D640A^ or Pol ε^DD875,877AA^.

Both Pol ε and Pol δ depend on the physical interaction with PCNA for processive DNA synthesis ^47^. We, therefore, tested if the Pol ε-PCNA interaction is important to compete Pol δ. To disrupt the Pol ε-PCNA interaction we purified a Pol ε^D640A-ΔPIP^ double-mutant complex, which in addition to the polymerase active site mutation contained the Pol2 FF1199,1200AA PIP-box mutation ^48^ (**Figure 1D**). Indeed, mutation of the PIP box of Pol2 attenuated the ability of Pol ε^D640A^ to compete Pol δ in both the primer extension assay (**Figure 3B**), and at replication forks (**Figure 3C**).

The N-terminal catalytic domain of Pol2 comprises a proofreading 3’-5’ exonuclease domain in addition to a polymerase domain ^7,49,50^. While the DNA polymerase activity of Pol2 normally dominates over the exonuclease, this balance is shifted towards the exonuclease under conditions that disfavor DNA synthesis, such as low dNTP concentrations or primer mismatches ^49,51^. Accordingly, we reasoned that mutation of the Pol2 polymerase active site may promote a switch to exonuclease activity, which may increase Pol2’s affinity for PCNA/primer-template junctions. We, therefore, purified a Pol ε complex in which essential active site residues in both the polymerase (D640) and exonuclease (D290/E292) domains were substituted with alanine (Pol ε^D640A-exo-^, **Figure 1D**) ^51^. Intriguingly, the ability of Pol ε^D640A-exo-^ to compete Pol δ in the primer extension assay was significantly reduced compared to Pol ε^D640A^, similar to that of Pol ε^D640A-ΔPIP^ (**Figure 3B**). Both inhibition of leading strand extension and excessive template unwinding at replication forks by Pol ε^D640A^ were also suppressed by the exonuclease mutation (**Figure 3C**). Thus, PCNA binding and a switch to exonuclease activity promote the competition of Pol δ by Pol ε harboring point mutations in the polymerase active site.

### Suppression of Pol δ inhibition by Pol ε restores viability of Pol2 polymerase point mutant cells

Next, we asked if mutations in the Pol2 PIP box suppress the lethality of catalytic point mutations in the Pol2 polymerase active site. Using a plasmid shuffle approach, we tested the ability of POL2 alleles expressed from an episomal vector to complement the deletion of chromosomal POL2. Since the inhibitory effect of catalytic point mutants of Pol ε on Pol δ was concentration-dependent (**Figures 2A** and **3B**), we expressed the episomal POL2 alleles from the GAL1,10 promoter to tune POL2 expression levels by modulating galactose concentrations in the medium. As shown in **Figure 4A**, this approach allowed us to vary the intracellular Pol2 concentration several-fold after transient growth in medium containing 0.05 % or 1 % galactose, respectively.

**Figure 4:**
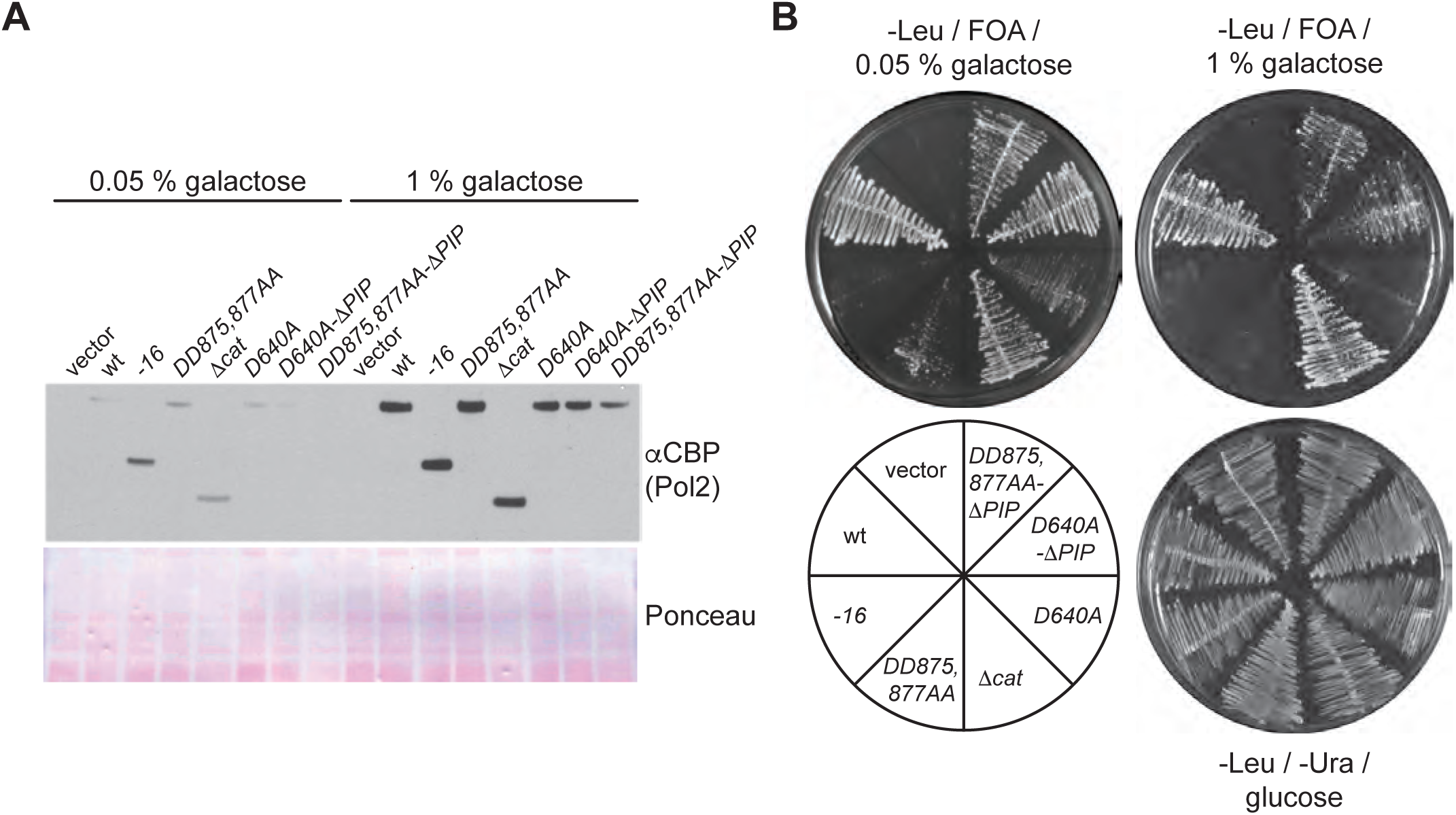
Inhibition of Pol ε’s capacity to compete Pol δ suppresses the lethality of Pol2 polymerase active site mutations. **(A)** Western blot analysis of galactose-induced Pol2 levels in cells expressing Pol2 variants from episomal plasmids. **(B)** POL2 plasmid shuffle assay. Tester cells were transformed with the indicated plasmids and assayed for growth on media containing glucose and lacking 5- FOA, or containing 5-FOA, 2 % raffinose, and the indicated concentrations of galactose.

While wild-type POL2 efficiently complemented the loss of endogenous POL2 at both 0.05 % and 1 % galactose, *pol2-16* was largely deficient in supporting cell growth at either galactose concentration (**Figure 4B**). As has been shown recently, growth of *pol2-16* cells is dependent on unidentified suppressors ^10^. It is possible that our plasmid-shuffle conditions do not provide sufficient time to accumulate suppressors prior to the plating of tester cells in 5-FOA. Unexpectedly, unlike *pol2-16*, *pol2Δcat* supported robust cell growth at both low and high galactose concentrations. This indicates that the reduced viability of *pol2-16* cells is not solely a consequence of the loss of the Pol2 catalytic domain. As expected, *pol2-D640A* and *pol2-DD875,877AA* did not support cell growth in the presence of 1 % galactose. However, simultaneous mutation of the PIP box in these alleles partially rescued cell growth at both 0.05 % and 1 % galactose. In general, the growth of all cells expressing *pol2* alleles with point mutations in the polymerase active site was enhanced on media containing low concentrations of galactose. These results demonstrate that reduced intracellular Pol2 levels and disruption of the interaction of Pol ε with PCNA/primer-template junctions suppress the lethality of point mutations in the Pol2 polymerase domain.

### Excessive DNA unwinding at replication forks upon dNTP depletion

To determine if uninterrupted helicase progression after polymerase inhibition is a general and intrinsic response of the replisome we sought to test additional conditions that impair polymerase activity during DNA replication. To this end, we monitored DNA replication *in vitro* under conditions of limiting dNTP concentrations. Incomplete leading strand extension to below half unit length was observed at dNTP concentrations of 0.25 μM and 0.5 μM, while close to full-length leading strands started to appear at 1 μM dNTPs (**Figure 5A**). In comparison, leading strands obtained in the presence of a high concentration (60 nM) of Pol ε^D640A^ ranged in length between those synthesized at 0.25 μM and 0.5 μM dNTPs. U* DNA was the predominant replication product at limiting dNTP concentrations, while regular θ-type replication intermediates formed the bulk of replication products at 1 μM dNTPs. This data demonstrates that polymerase inactivation by dNTP depletion, similar to mutational inactivation of Pol ε, causes excessive DNA unwinding at the budding yeast replication fork.

**Figure 5:**
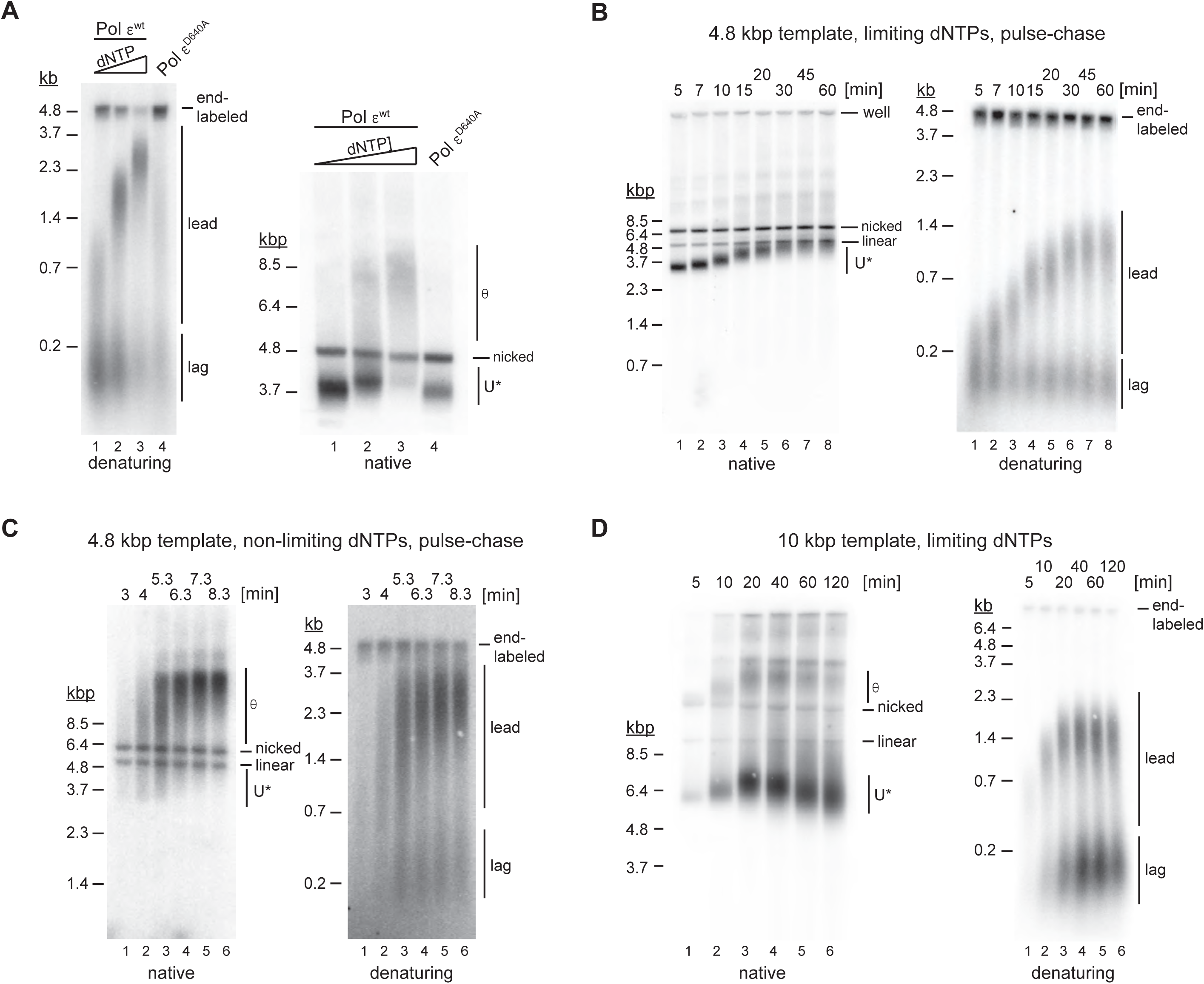
Helicase-polymerase uncoupling at limiting dNTP concentrations. **(A)** Standard replication reactions were performed in the presence of 0.25 μM, 0.5 μM, or 1 μM dNTPs and 30 nM Pol ε^wt^ (lanes 1-3), or 40 μM dNTPs and 60 nM Pol ε^D640A^ (lane 4). Template: pARS1. **(B)** Pulse-chase experiment of standard replication reaction performed in presence of 0.25 μM each dNTP. Pol ε^exo-^ was used in place of Pol ε^wt^. Times indicate minutes after origin activation. Chase (500 μM cold dATP) was added at 5 minutes. Template: pARS1. **(C)** Pulse-chase experiment of standard replication reaction at non-limiting dNTP concentrations (40 μM dCTP / dTTP / dGTP, 4 μM dATP). Times indicate minutes after origin activation. Chase (500 μM cold dATP) was added at 5 minutes. Template: pARS1. **(D)** Time course analysis of standard DNA replication reaction performed at 0.25 μM dNTPs (each); Template: pARS305.

We sought to exclude the possibility that resection of nascent strands upon dNTP exhaustion could be the source for unwound DNA in U*. This possibility arises from the fact that limiting dNTP concentrations shift the balance between the DNA polymerase and 3’-5’ exonuclease activities of Pol ε toward exonuclease activity ^51^. To characterize the kinetics of U* formation under limiting dNTP concentrations we performed pulse-chase experiments in the presence of exonuclease-deficient Pol ε. For this, DNA synthesis was initiated in the presence of 0.25 μM dNTPs, including α-^32^P-dATP, and chased with 500 μM cold dATP 5 minutes after origin activation, which maintains the remaining three dNTPs at limiting concentrations throughout the chase. As shown in **Figure 5B**, U* DNA of maximal gel mobility was formed early in the reaction, while the gel mobility of U* molecules subsequently continued to decrease as DNA synthesis progressed. The rate of leading strand synthesis was reduced by an order of magnitude, progressing at a maximum rate of 55 bp/min during the first 15 minutes of the reaction and decreasing at later time points, before stalling at a length below the normal half-unit length. Similar results were also obtained in the presence of wild-type Pol ε (**Figure S8**), demonstrating that the formation of U* under limiting dNTP conditions is also not a consequence of increased strand-displacement by Pol ε^exo- 52^. Moreover, we note that U* was formed under limiting dNTP conditions irrespective of the presence of Mrc1 and Csm3-Tof1 (**Figure S8**), which suppress the progression of replisomes from sites of DNA synthesis in HU-treated cells ^25^. Our data indicate that Mrc1/Csm3-Tof1 do not act downstream of checkpoint activation to restrain replisomes at sites of ongoing DNA synthesis. In summary, we conclude that the initial formation of U* under limiting dNTP conditions is due to excessive DNA unwinding by CMG, and not 3’-5’ resection of nascent strands.

By comparison, in pulse-chase experiments under non-limiting dNTP conditions we find not only that the rate of DNA synthesis is greatly increased, but also that the bulk of replication intermediates progresses, as expected, through canonical θ intermediate stages (**Figure 5C**). Importantly, U* DNA is transiently detectable at early time points (3-5 minutes after origin activation) even under non-limiting dNTP conditions, indicating that the formation of U* is not dependent on the inhibition of DNA polymerases by dNTP depletion.

The decrease of the gel mobility of U* concomitant with DNA synthesis in the above experiments suggested that DNA synthesis and DNA unwinding under limiting dNTP conditions occurred at comparable rates (**Figures 5B** and **S8**). However, analyses over longer time periods and on larger plasmid templates revealed that the gel mobility of U*, after initially decreasing with ongoing DNA synthesis, continued to increase again once DNA synthesis had stalled, indicating that DNA unwinding by CMG continued upon dNTP exhaustion (**Figure 5D**). Thus, formation of U* under limiting dNTP conditions results from uninterrupted and continuous helicase progression uncoupled from DNA synthesis.

### Rad53 inhibits excessive DNA unwinding at replication forks after polymerase inhibition

Previous studies had suggested that the replication checkpoint kinase, Rad53, inhibits replisome progression in yeast cells treated with HU ^24,25,32^. We, therefore, tested the ability of purified recombinant Rad53 to inhibit CMG at reconstituted forks. Because Rad53 also inhibits origin firing ^53–55^, we used a two-step approach to temporally separate fork progression from origin firing *in vitro* (**Figure 6A**). In the first step, replication forks were transiently stalled by omission of topoisomerase from the origin activation reaction. In the second step, the topological block to fork progression was resolved by addition of Top1 and Top2, while concomitant DNA synthesis was controlled with aphidicolin.

**Figure 6:**
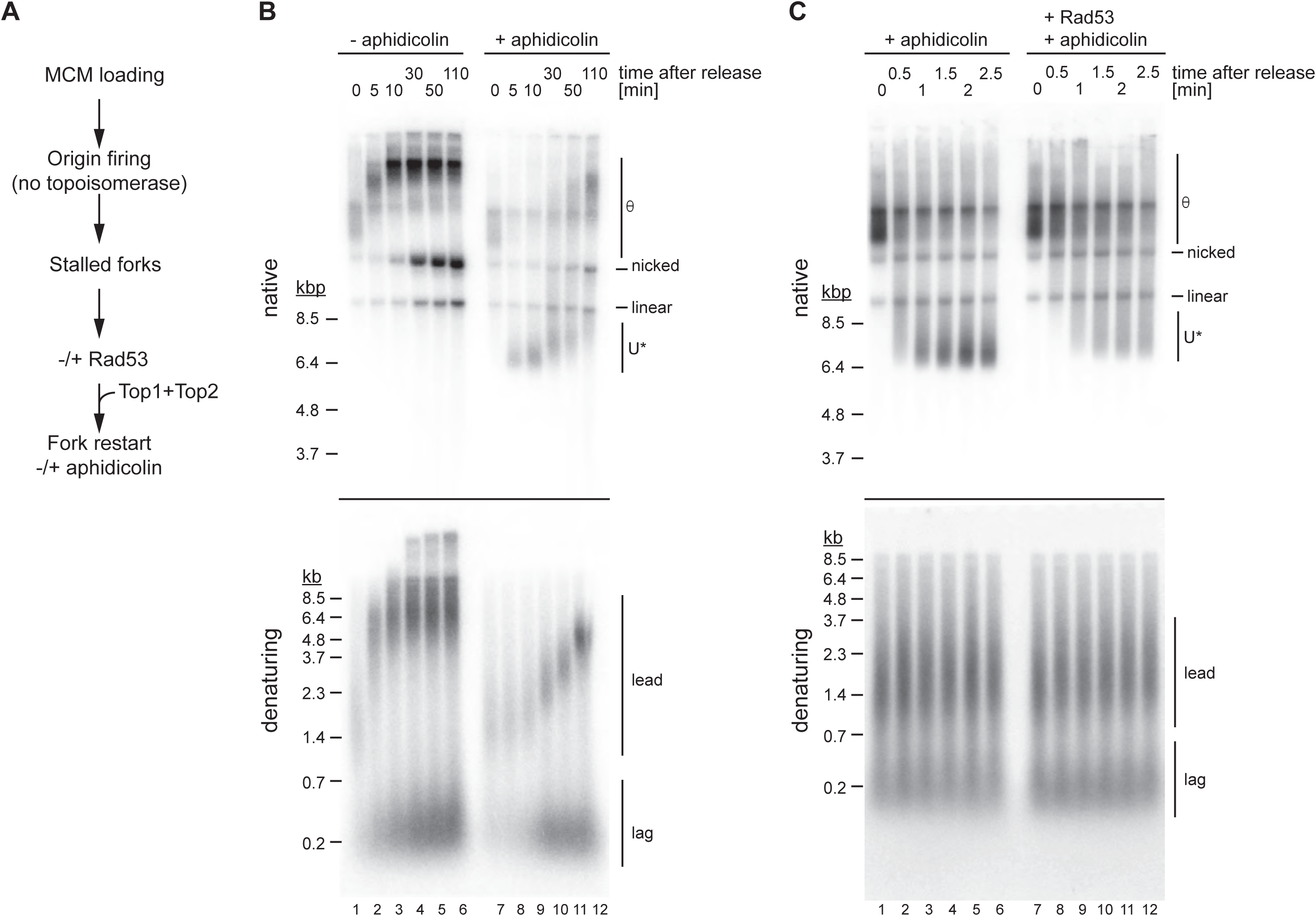
Rad53 inhibits DNA unwinding after helicase-polymerase uncoupling in presence of aphidicolin. (**A**) Experimental outline. (**B**) Effect of aphidicolin on fork progression. Forks were released from a topological stall in the absence or presence aphidicolin (30 μM) as indicated. Times indicate minutes after Top1/Top2 addition. Template: pARS305. (**C**) Effect of Rad53 on helicase-polymerase uncoupling. Stalled forks were either mock-treated (lanes 1-6) or treated with Rad53 (lanes 7-12) prior to release in the presence of aphidicolin. Template: pARS305.

**Figure 6B** shows that on the 10 kbp plasmid template replication forks stalled approximately 1-2 kb from the origin in the absence of topoisomerase, resulting in the formation of early θ replication intermediates. Addition of Top1 and Top2 without aphidicolin led to efficient fork release, resulting in the formation of fully extended leading strands around 10 minutes after fork release, progression of the θ replication intermediates, and formation of decatenated plasmid daughters within 30 minutes of fork restart. In contrast, in the presence of aphidicolin, early θ replication intermediates were rapidly converted to U* DNA and DNA synthesis progressed at a greatly reduced rate. Consistent with U* being a negatively supercoiled product of DNA unwinding, U* formed in the presence of aphidicolin is efficiently relaxed by *E. coli* Topo I (**Figure S9**). Thus, inhibition of DNA polymerase activity by aphidicolin, like mutational inactivation of Pol ε or dNTP depletion, results in apparently immediate and excessive DNA unwinding at the replication fork *in vitro*.

As maximal U* mobility was reached within 5 minutes after fork release in the presence of aphidicolin (**Figure 6B**), we next assessed the effect of purified recombinant Rad53 on the kinetics of U* formation at early time points within a 2.5 minute window after fork release in aphidicolin; DNA synthesis did not progress detectably within this period under these conditions (**Figures 6C**). In the absence of Rad53, U* of maximal gel mobility was efficiently formed 1-1.5 minutes after fork release. However, both the rate of U* formation and the final extent of unwinding of U* were reduced in the presence of Rad53. The inhibition of DNA unwinding at replication forks in the presence of aphidicolin requires the kinase activity of Rad53, as U* formation was not inhibited in the presence of kinase-dead Rad53 (**Figure S10**). Thus, Rad53 controls DNA unwinding at replication forks following polymerase inhibition.

### Rad53 control of the CMG helicase is not dependent on DNA polymerase inhibition

To identify potential targets of Rad53 in the replisome that mediate the inhibition of DNA unwinding after polymerase inhibition we initially focused on Mrc1 and Csm3-Tof1. Both proteins have been shown to promote fork rates ^13,56,57^, are phosphorylated by Rad53 during replication stress ^58,59^, and are required for the restriction of replisome progression in cells treated with HU ^25^. However, using the approach of **Figure 6C**, we find that Rad53 can limit DNA unwinding after polymerase inhibition also in the absence of Csm3-Tof1 and Mrc1(**Figure 7A**).

**Figure 7:**
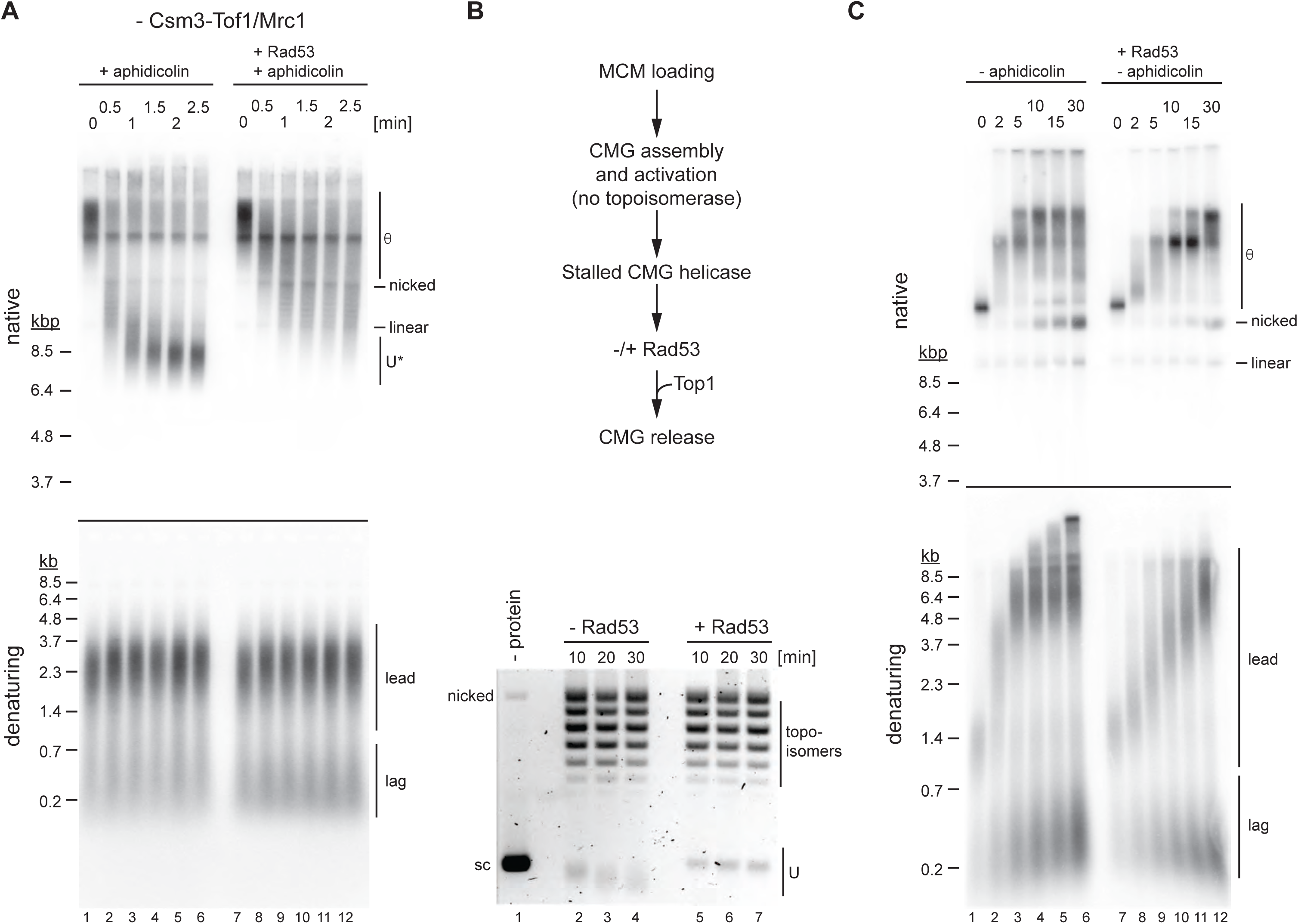
Rad53 controls the CMG helicase to limit DNA unwinding after helicase-polymerase uncoupling. (**A**) As Figure 6B, except that Csm3-Tof1 and Mrc1 were omitted from the reaction. Template: pARS305. (**B**) CMG helicase assay. Top: Reaction scheme. Bottom: Products analyzed by ethidium-bromide stain after native agarose gel-electrophoresis. Times indicate minutes after Top1 addition. sc: supercoiled. Template: pARS1. (**C**) As outlined in Figure 6A, omitting aphidicolin and including Rad53 as indicated. Template: pARS305.

Next, we considered the possibility that the CMG helicase itself is regulated by Rad53. To test this hypothesis, we modified a previously reported approach that allows the assembly and activation of the CMG helicase with a limited set of initiation factors that avoids the assembly of complete replisomes (**Figure 7B**) ^45^. To separate the inhibitory effect of Rad53 on CMG assembly from its potential role in regulating the CMG helicase, we assembled and activated CMGs on plasmid DNA in the absence of topoisomerase to stall active CMGs at template supercoils. Stalled CMGs were mock-treated or treated with Rad53, before being released from the topological block by addition of Top1. CMG helicase activity is detected by the generation of U form DNA in native gels. While the gel-mobility of U form DNA generated prior to CMG stalling continued to increase after block removal in the absence of Rad53, it persisted in the presence of Rad53, demonstrating that Rad53 inhibits DNA unwinding by CMG. We note that this data does not exclude the possibility that proteins other than CMG are targeted by Rad53 to regulate CMG activity, as replisome components such as Pol ε and Mcm10 are required to assemble and activate CMG at origins and are, therefore, also present in the CMG helicase assay here.

To determine if Rad53 limits CMG activity only after inhibition of DNA polymerases, we employed the approach of **Figure 6A** to test the ability of Rad53 to inhibit fork progression in the absence of aphidicolin (**Figure 7C**). As before, in the absence of Rad53, addition of Top1 and Top2 induced efficient and synchronous release of forks stalled by topological strain 1-2 kb from the origin, with leading strand synthesis largely complete around 5 minutes after fork release, and fully replicated and segregated plasmid daughters starting to appear between 5-10 minutes after fork restart. In contrast, fork progression was markedly reduced when forks were incubated with Rad53 during and after release from the block, with fully replicated products not apparent before 30 minutes after fork release. Thus, Rad53 inhibition of fork progression is not dependent on the inhibition of DNA polymerases by aphidicolin.

## DISCUSSION

We have shown that inhibition of DNA polymerase activity at a eukaryotic replication fork by mutational inactivation of Pol ε, dNTP depletion, or chemical inhibition causes uninterrupted CMG helicase progression uncoupled from DNA synthesis. This indicates an absence of replisome-intrinsic mechanisms that coordinate the activities of the CMG helicase and DNA polymerases at the fork. While under normal conditions the inherent rates of the CMG helicase and DNA polymerases may balance DNA unwinding and synthesis at the fork, the lack of intrinsic control mechanisms would explain the dependency on checkpoint pathways to maintain fork integrity during replication stress that creates an imbalance between helicase and polymerase activities. Moreover, this raises the possibility that helicase-polymerase uncoupling occurs stochastically at eukaryotic forks, i.e. DNA synthesis and DNA unwinding are not tightly coupled at the eukaryotic replisome. Consistent with this notion, we observe U* form DNA also under normal replication conditions (**Figure 5C**). Combined with the observation that lagging strand synthesis can also occur uncoupled from leading strand synthesis at eukaryotic forks ^22,24^, this may indicate stochastic features of the eukaryotic replisome analogous to those observed in *E. coli*, which has been proposed to increase the robustness of the DNA replication process ^60^.

Recoupling of the CMG to leading strand synthesis after stochastic or stress-induced uncoupling is likely to involve Pol δ, which exhibits a rate of DNA synthesis on RPA-coated ssDNA that exceeds normal fork rates ^12,13,61^, and which competes free Pol ε at PCNA/primer-template junctions ^12,14^. This recoupling activity of Pol δ, in addition to its role in initiating the leading strand ^10,62,63^, may thus contribute to Pol δ activity on the leading strand ^64^.

Stochastic helicase-polymerase uncoupling increases the risk of ssDNA exposure at the fork. In *E. coli*, the intrinsic slowing of the replicative helicase upon uncoupling from the polymerase has been likened to a “dead-man’s switch” that limits the generation of ssDNA and facilitates the recoupling of helicase and polymerase ^60^. Some evidence suggests that CMG inherently unwinds DNA also at rates that are below those of normal forks ^19–21^, which may be due to CMG slippage, loss of the driving force of DNA synthesis, or a switch from ssDNA to dsDNA binding ^18,19,65,66^. Consistent with this notion, our data suggests that the rate of DNA unwinding by yeast CMG does not exceed ∼ 55 bp per minute if uncoupled from DNA synthesis (**Figure 5B**), well below the normal fork rate of ∼ 1-2 kb per minute. Thus, as in bacteria, intrinsic helicase slowing upon uncoupling from DNA synthesis may act as a first response to buffer helicase-polymerase uncoupling at eukaryotic forks.

However, intrinsic slowing of CMG is not sufficient to maintain fork function during replication stress, as forks stalled by HU treatment in *rad53Δ* cells exhibit longer tracts of ssDNA than forks in wild-type cells and are restart defective ^24,36^. Rad53 restricts the progression of replisomes away from sites of DNA synthesis, which limits the exposure of ssDNA at stalled forks ^24,25,32^. Our demonstration that Rad53 inhibits the rate of DNA unwinding by CMG thus suggests a molecular mechanism for this checkpoint function. Restriction of replisome progression by Rad53 may promote fork stability by limiting the generation of ssDNA to prevent replication catastrophe from RPA exhaustion and protect forks from nuclease attack ^35,67,68^. CMG inhibition by Rad53 may also help maintain the replisome near the site of polymerase stalling to facilitate the recoupling of CMG to leading strand synthesis. Alternatively, reducing the amount of unwound parental DNA may promote fork restart by fork reversal mechanisms that may be limited by the processivity of the strand annealing activity of fork remodeling enzymes ^69^.

How Rad53 inhibits the CMG helicase is not known. Known Rad53 targets in the replisome include Mrc1, Tof1, Ctf4, and Cdc45 ^58,59,70^. While Mrc1 activates Rad53 during replication stress, our data demonstrate that Rad53 can inhibit DNA unwinding by CMG also in the absence of Mrc1 and Tof1. Phosphorylation of Cdc45 creates a docking site for Rad53 in the replisome, but it is not known if this interaction regulates CMG progression or mediates some other aspect of Rad53 function ^70^. The assays developed here will allow to interrogate this question using conditions that specifically disrupt the Cdc45-Rad53 interaction. The inhibition of CMG activity by Rad53 may also involve Mcm10 or Pol ε, both of which have been implicated in modulating CMG activity and bind CMG in proximity to Rad53 ^6,71–73^. Mcm2-7 subunits are targeted by a number of kinases that regulate DNA replication, including Mec1/ATR, DDK, and CDK, while GINS subunits have been identified as targets for Mec1 in yeast and Chk2 in *Drosophila*, raising their potential as targets for Rad53 ^1,32,34, 74–76^. Indeed, Chk2 has been shown to inhibit the helicase activity of recombinant *Drosophila* CMG in the absence of other replisome components, suggesting that CMG phosphorylation can directly modulate CMG activity ^74^.

The inhibition of CMG activity by Rad53 does not require inhibition of DNA polymerase activity. Nontheless, it is possible that Rad53 limits DNA unwinding specifically after stochastic CMG uncoupling from DNA synthesis. Alternatively, Rad53 may inhibit CMG irrespective of its coupling to DNA synthesis. This would support a model in which stressed forks that have generated sufficient ssDNA for Rad53 activation in the wake of CMG may also inhibit normal fork progression in *trans* (**Figure S11A**). Such a mechanism may explain the global slowing of replication forks in yeast cells treated with HU and provide an additional strategy to help preserve dNTP levels during replication stress ^28^. While Mec1-Ddc2-dependent Rad53 activation is bypassed here by the use of autophosphorylated recombinant Rad53 ^77^, reconstitution of this pathway will provide a gateway to differentiate between global and local Rad53 effects in the future.

Our observation that point mutations in the Pol ε polymerase domain cause CMG uncoupling from DNA synthesis was unexpected. While an inactive polymerase domain may compete other polymerases from primer ends, our finding that free mutant Pol ε complexes compete Pol δ at PCNA/primer-template junctions was unexpected given the greater affinity of Pol δ for PCNA/primer-template junctions ^14^. However, inhibition of the Pol2 polymerase and the concomitant switch to exonuclease activity may increase the affinity of Pol ε for PCNA-bound primer ends, allowing it to compete Pol δ off PCNA/primer-template junctions. The upregulation of dNTP levels by the checkpoint may thus limit Pol ε’s potential to compete other polymerases involved in replication restart.

At limiting concentrations of Pol ε polymerase point mutants, leading strand synthesis by Pol δ proceeds at a rate equivalent to that observed in the presence of Pol ε^Δcat^, indicating a similar mode of fork progression (**Figure S11B, ii** and **iv**). However, at concentrations that exceed the number of active forks, free mutant Pol ε complexes compete with Pol δ, resulting in a reduction of the rate of leading strand synthesis (**Figure S11B, iii**). Our data indicate that this inhibition is the cause of lethality by *pol2* alleles harboring an inactive polymerase domain. However, it has been estimated that most or all intracellular Pol ε is bound to CMG ^14^. This estimate was based on proteomic data indicating similar intracellular concentrations of Pol ε and CMG subunits, and the assumption that all CMG subunits in the cell are assembled into CMG complexes. However, only a fraction of intracellular MCM subunits is loaded onto chromatin in G1 phase ^78,79^, and only a fraction of chromatin-loaded MCMs is converted to CMGs at any one time ^1^, indicating that the majority of Pol ε is not constrained at CMGs, consistent with our model. We note that Pol δ and Pol ε are also present at comparable concentrations in the cell ^80^. Since we observe significant competition of Pol δ at equal concentration of polymerase-dead Pol ε *in vitro*, this data is consistent with our model that the lethality of POL2 polymerase point mutants is due to Pol δ inhibition.

Contrary to *pol2-16,* we find here that the *pol2Δcat* allele and polymerase-dead *pol2* alleles lacking a functional PIP box efficiently rescue the lethality of POL2 deletion. Thus, the loss of leading strand synthesis by Pol ε is significantly better tolerated by cells than previously thought, although it remains to be seen how efficient and stress-resistant replication forks are in cells lacking Pol ε polymerase activity. The enhanced sickness of *pol2-16* compared to *pol2Δcat* cells is, therefore, likely associated with the additional 301 amino acid residues at the *pol2-16* N-terminus (**Figure 1C**). This region encompasses the PIP box, which may allow *pol2-16* to compete other DNA polymerases off PCNA/primer-template junctions, or it may decrease the stability of *pol2-16* ^10^. In any case, plasticity in both polymerase usage and helicase-polymerase coupling are likely to contribute to the robustness of the DNA replication process in eukaryotic cells.

## DATA AVAILABILITY

Source data for graphs in Figures 2B, 2D, 3B, S2A, and S3 are available in Supplementary Table 1. All other data are available upon request.

## ACKNOWLEDGMENTS

This work was supported by NIH grant R01GM107239. We thank Ken Marians and Tom Kelly for critical comments on the manuscript, and Xiaolan Zhao for yeast strains.

## AUTHOR CONTRIBUTIONS

D.R. and S.D. conceived this study and designed the experiments. S.D. performed the experiments. D.R. wrote the paper with help from S.D.

---

**Figure S1: Related to Figure 1.**
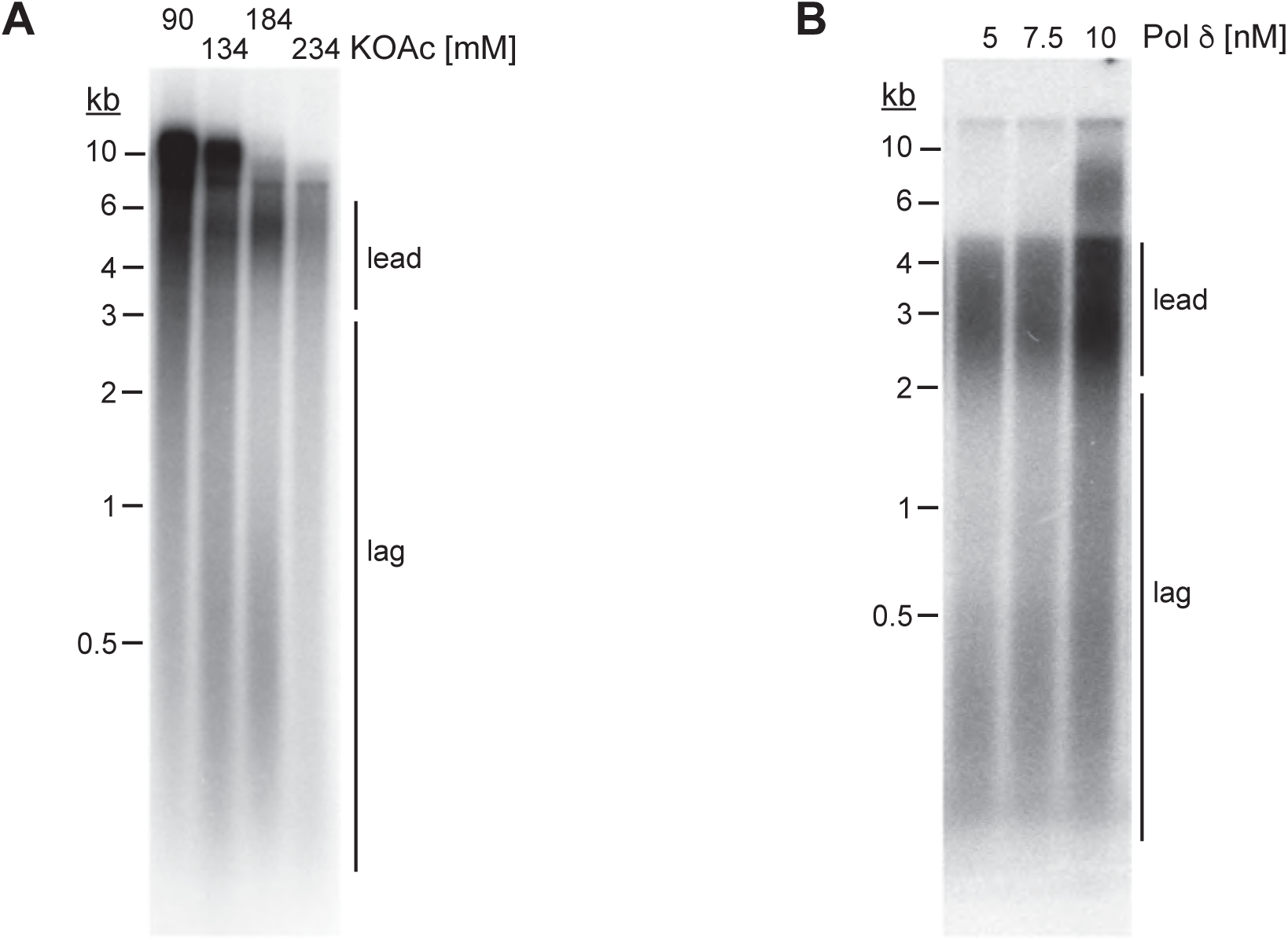
Reaction conditions affecting lagging strand synthesis *in vitro*. **(A)** Standard replication reactions were performed at various final salt concentrations as indicated. Template: pARS305. **(B)** Titration of Pol δ into standard replication reactions. Template: pARS1. Reaction products were analyzed by 0.8 % alkaline agarose gel-electrophoresis and autoradiography.

**Figure S2: Related to Figure 2.**
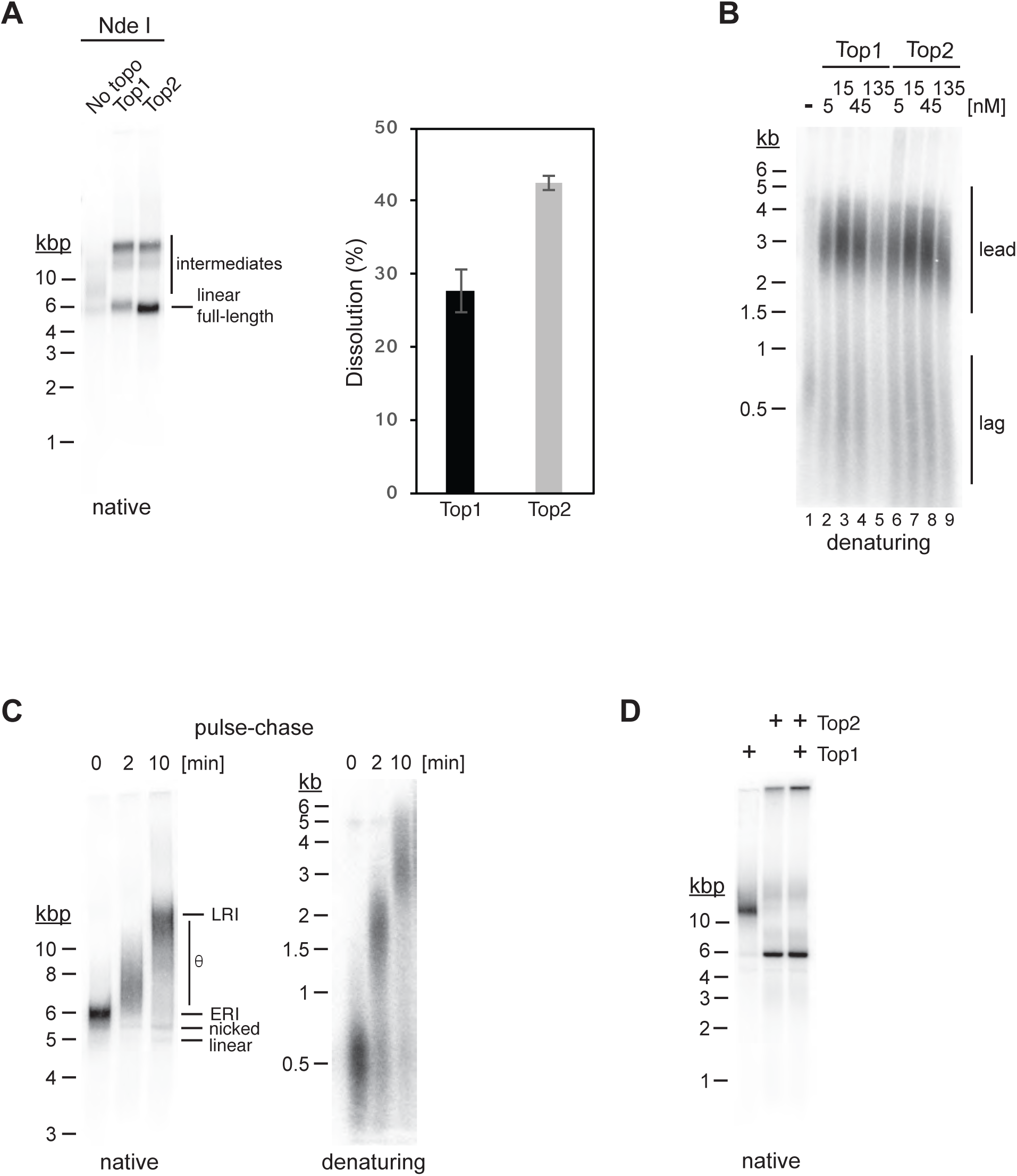
Characterization of replication products. **(A)** Standard replication reactions with pARS1 were carried out in the absence or presence of Top1 or Top2, as indicated. Reactions were stopped 60 minutes after origin activation, and replication products analyzed by native agarose gel-electrophoresis and autoradiography after linearization with the unique cutter Nde I, which cuts near the origin (fully replicated DNA molecules will resolve into linear monomers after linearization, whereas replication products containing unreplicated regions will resolve into double Y-shaped intermediates. Representative gel is shown on the left. The bar diagram depicts the average dissolution efficiency (fraction of linear full-length molecules per reaction) and standard deviation of three independent experiments. **(B)** Replication products from experiment in Figure 1B were analyzed by alkaline agarose gel-electrophoresis and autoradiography. **(C)** Replisomes were formed on chromatin templates in the presence of α-^32^P-dCTP and stalled by omission of topoisomerase from the reaction (lane 1). After 15 minutes, Top1 was added to the reaction to release the stalled replisomes and intermediates chased by simultaneous addition of excess cold dCTP. At the indicated time points (lanes 2 and 3) replication products were isolated and analyzed by native (left) or denaturing agarose gel-electrophoresis and autoradiography. (**D**) Standard replication reactions carried out in the presence of either Top1, Top2, or both. Replication products were analyzed by native agarose gel-electrophoresis and autoradiography.

**Figure S3: Related to Figure 1.**
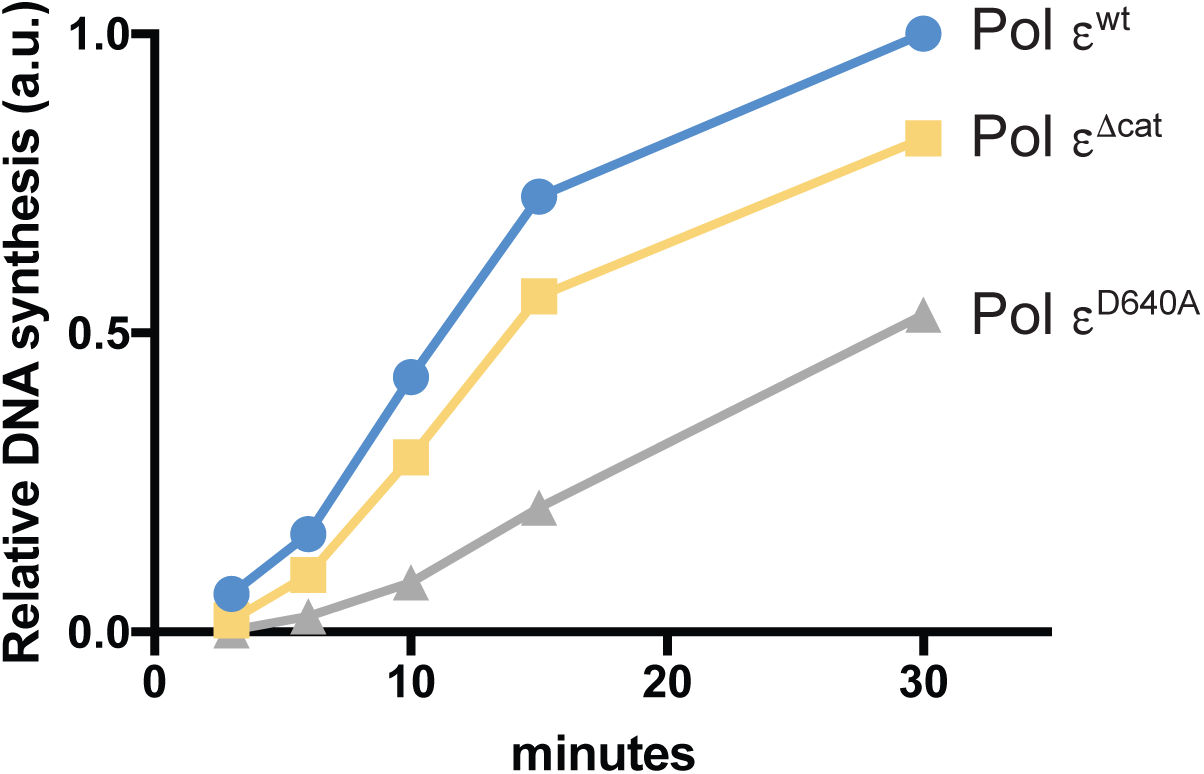
Reduced DNA synthesis in the presence of Pol ε polymerase mutants. Total relative DNA synthesis in reactions of Figure 1E were measured using ImageJ and plotted over time.

**Figure S4: Related to Figure 1.**
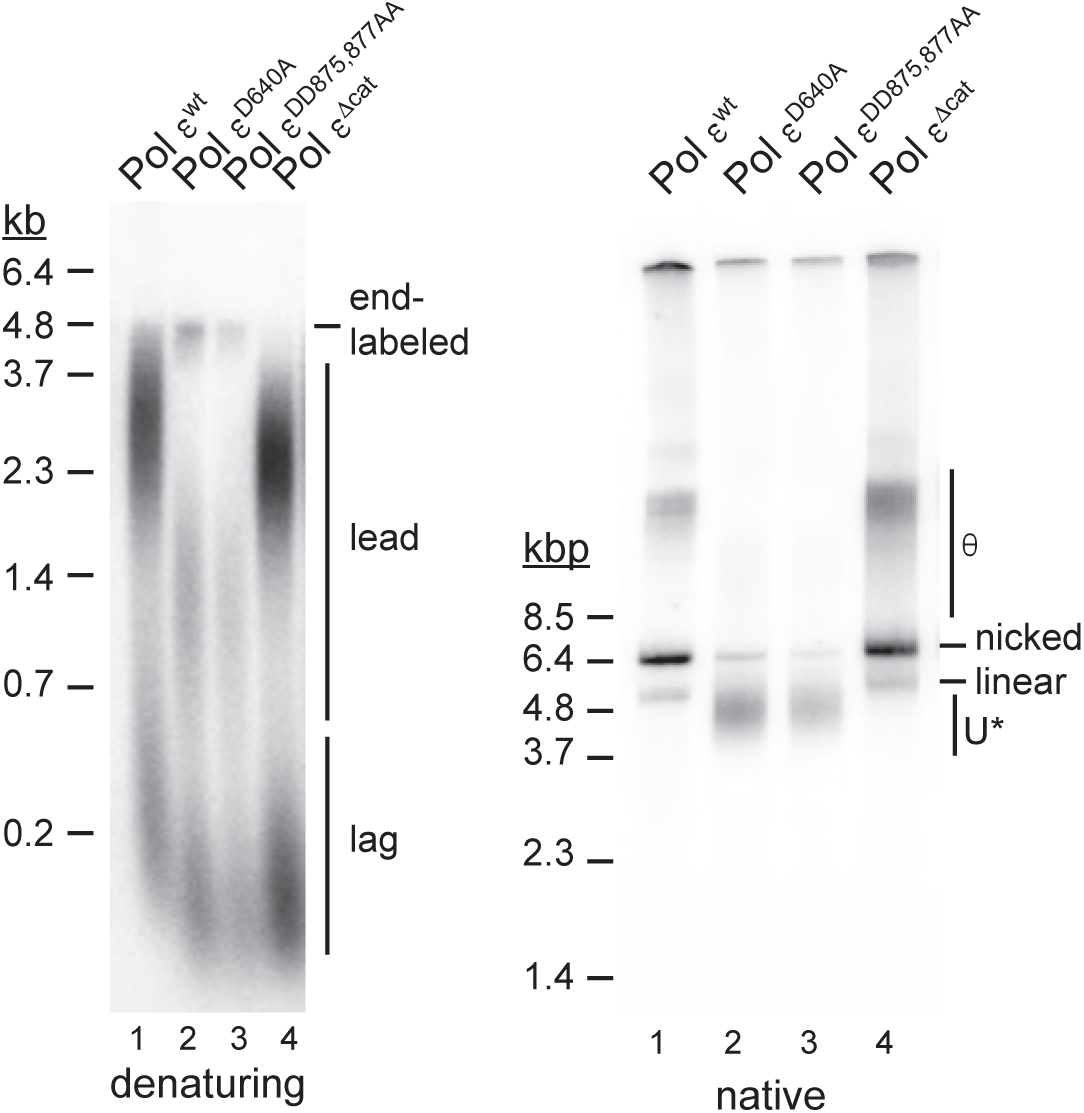
Inhibition of fork progression by Pol ε polymerase mutant complexes. Standard DNA replication reactions were carried in the presence of wild-type or the indicated mutant Pol ε variants (60 nM). 45 minutes after origin activation reactions were stopped and replication products analyzed by alkaline (left) or native (right) agarose gel-electrophoresis and autoradiography. Template: pARS1.

**Figure S5: Related to Figure 1.**
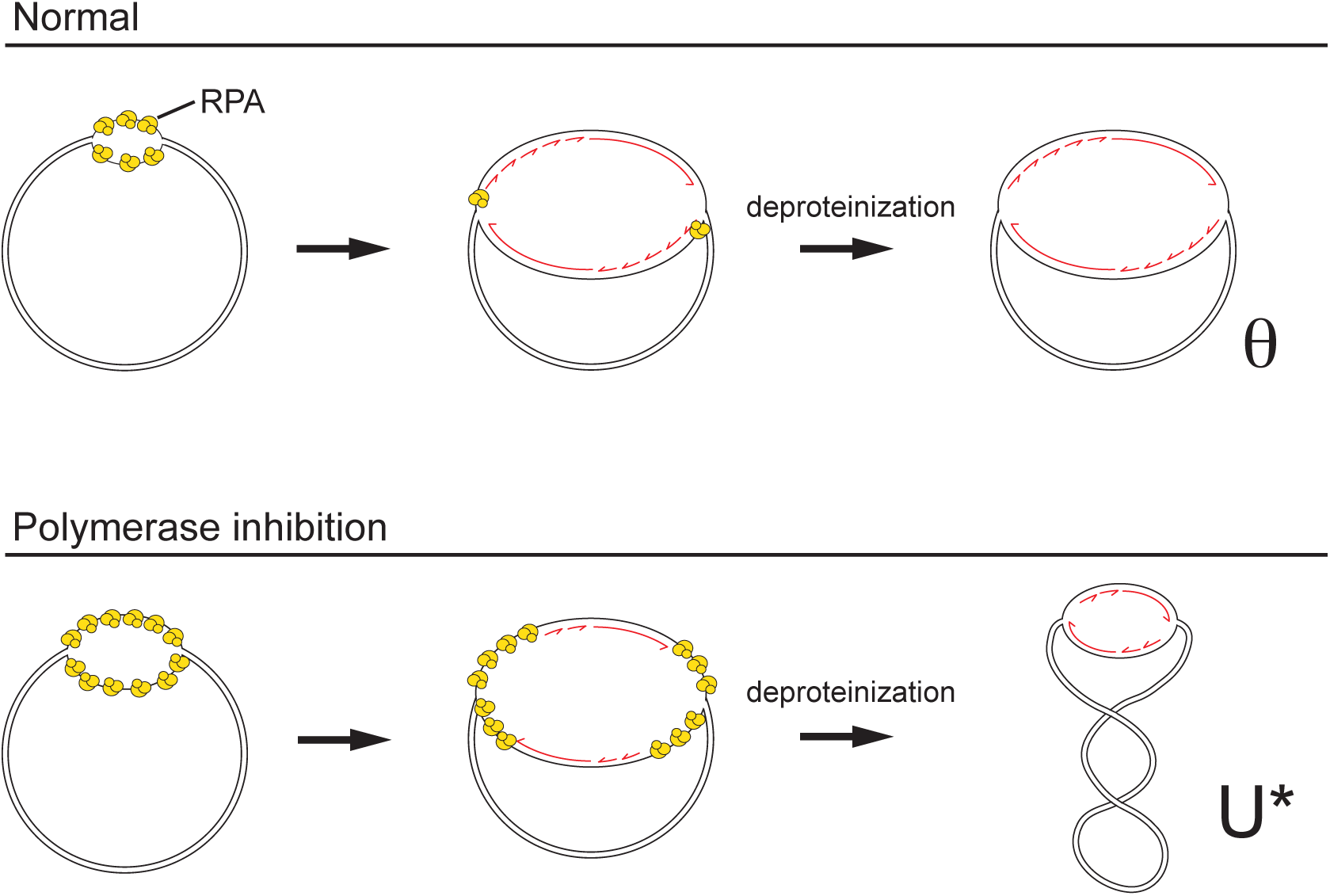
Model for formation of θ and U* replication intermediates during plasmid replication *in vitro*. In normal DNA replication, the origin is initially unwound upon CMG activation (top left), followed shortly thereafter by the commencement of DNA synthesis and the coupling of leading strand synthesis to DNA unwinding by CMG (top center). Compensatory positive supercoils formed in the template during unwinding and fork progression are removed by Top1 and/or Top2. After deproteinization, the resulting θ structure is maintained (top right). In contrast, under conditions that slow-down DNA synthesis after origin unwinding the CMG helicase progresses along the template (bottom left) in advance of DNA synthesis; compensatory positive supercoils generated during DNA unwinding are removed by Top1 and/or Top2, and the unwound single-stranded DNA is stabilized by RPA binding (bottom center). Upon deproteinization, unwound complementary DNA strands reanneal, causing compensatory negative supercoils and thus resulting in a partially replicated, negatively supercoiled replication intermediate, U* (bottom right).

**Figure S6: Related to Figure 2.**
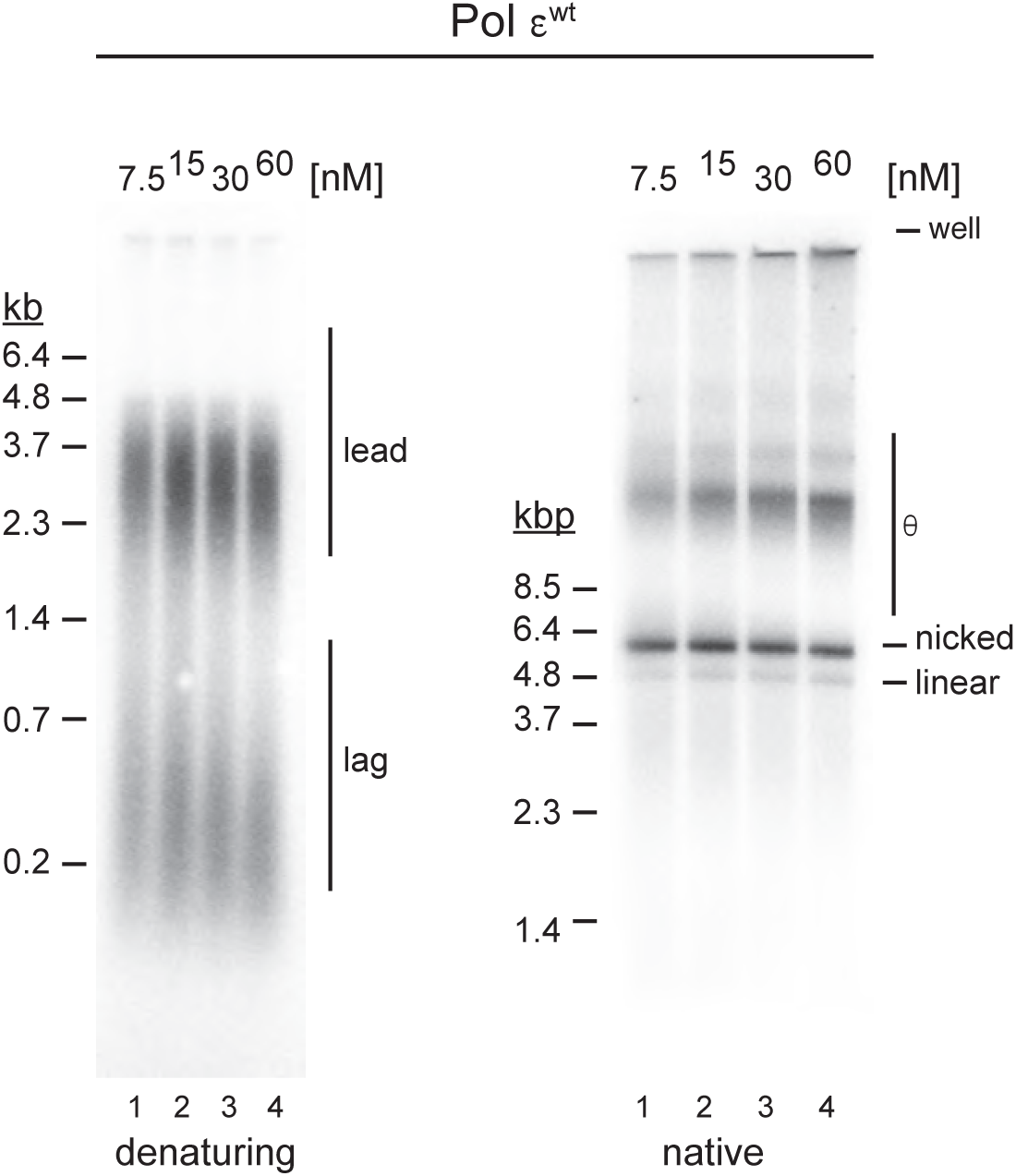
Effect of Pol ε concentration on fork progression. Pol ε^wt^ was titrated into standard replication reactions, reactions stopped 45 minutes after origin activation, and replication products analyzed by denaturing (left) or native (right) agarose gel-electrophoresis and autoradiography as indicated. Template: pARS1.

**Figure S7: Related to Figure 3.**
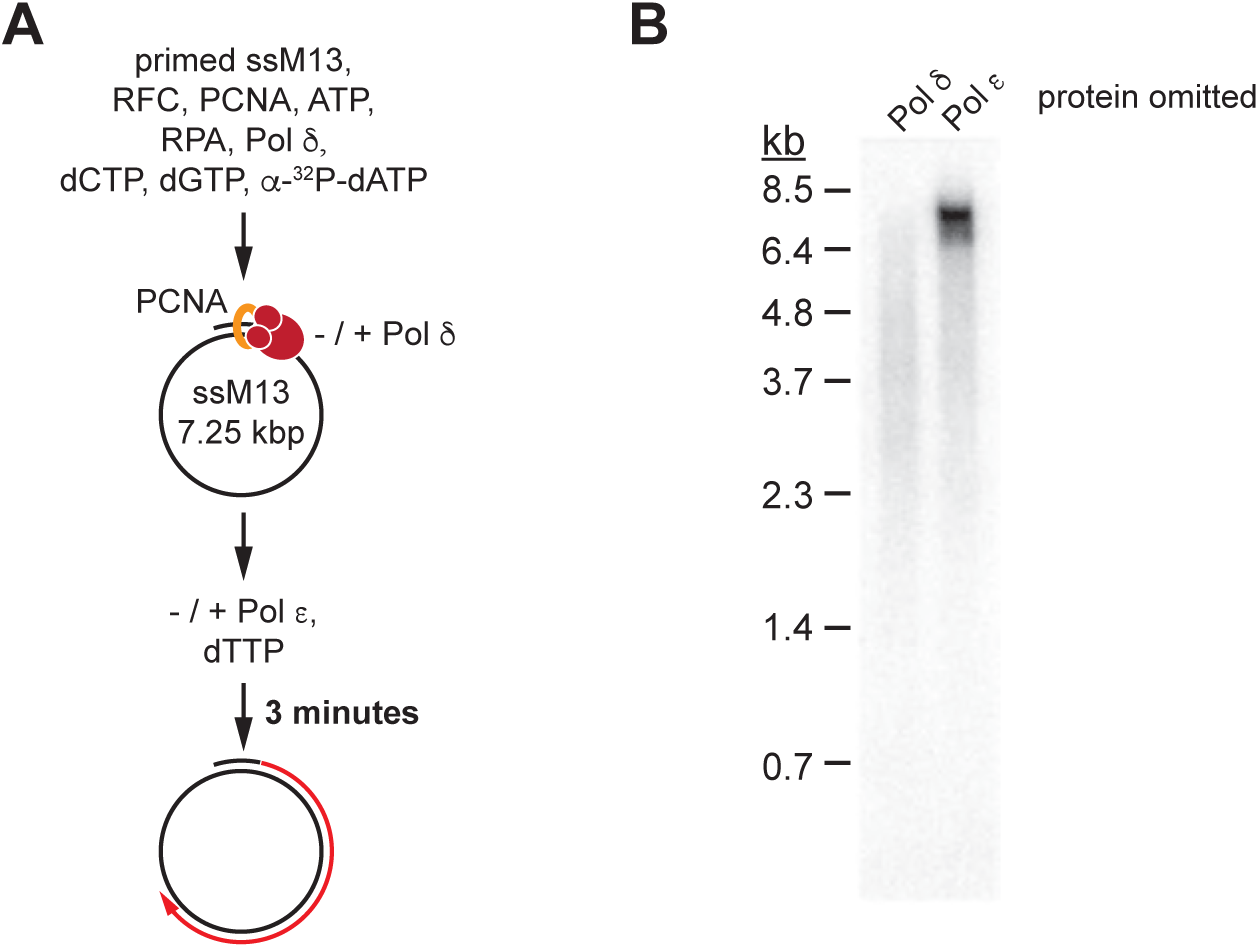
Primer extension products obtained with Pol δ or Pol ε. **(A)** Reaction scheme: Singly primed single-stranded M13mp18 DNA was pre-incubated with RPA, RFC/PCNA, three nucleotides, and Pol δ to initiate primer extension; Pol ε was subsequently added along with the remaining fourth nucleotide and incubation continued for 3 minutes. **(B)** Denaturing agarose gel analysis of primer extension products obtained according to reaction scheme in (A), but with either Pol δ or Pol ε omitted from the reaction.

**Figure S8: Related to Figure 5.**
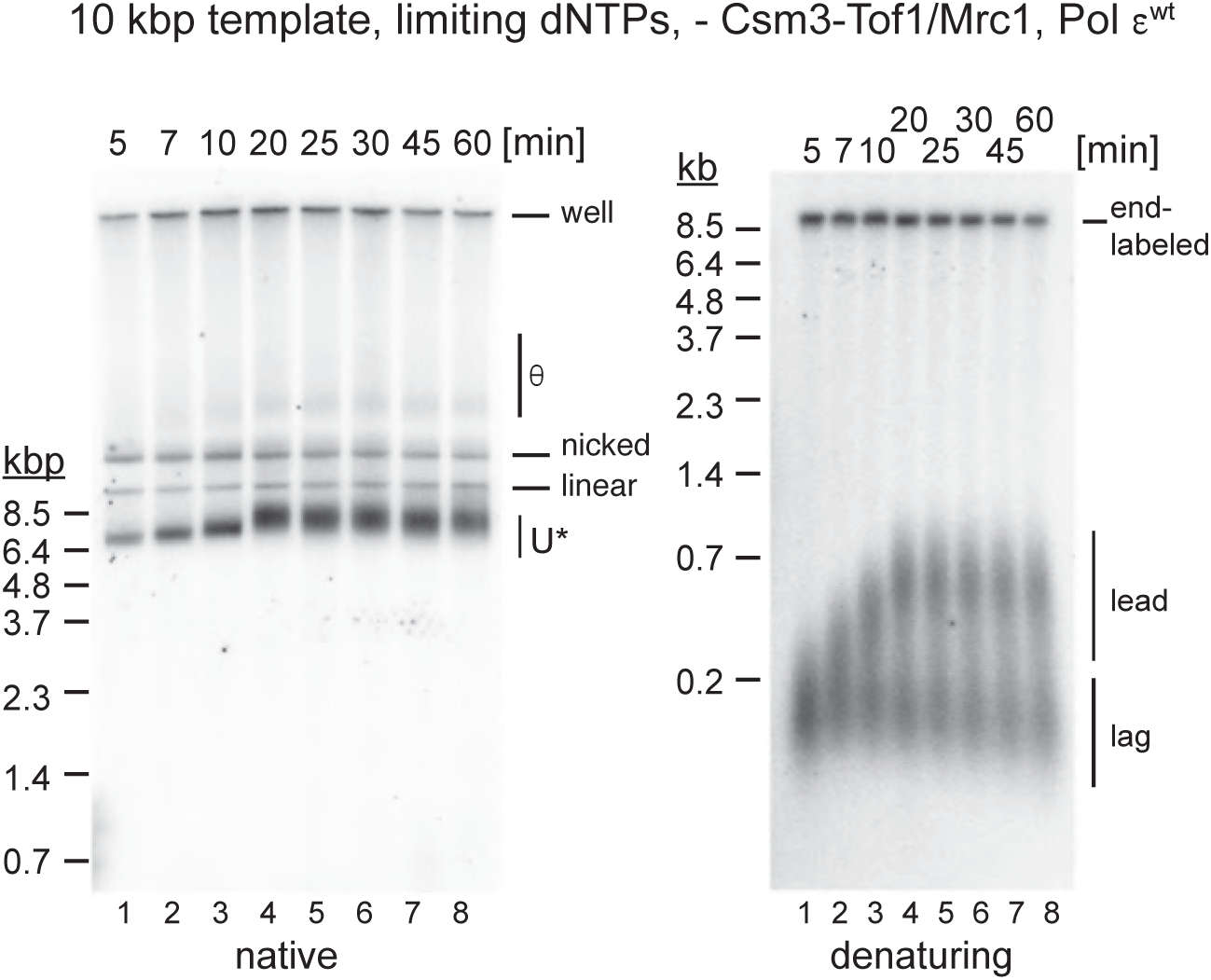
Excessive DNA unwinding under limiting dNTP conditions in the absence of Csm3-Tof1/Mrc1 and presence of Pol ε^wt^. Pulse-chase experiment of standard replication reaction performed at 0.25 μM each dNTP as in Figure 5B, with the following changes: 1) Pol ε^wt^ was used instead of Pol ε^exo-^; 2) Csm3-Tof1 and Mrc1 were omitted from the reaction; 3) pARS305 instead of pARS1 served as a template. Time indicates minutes after origin activation. The reaction was chased with 500 μM cold dATP 5 minutes after origin activation. Reaction products were analyzed by denaturing agarose gel-electrophoresis and autoradiography.

**Figure S9: Related to Figure 6.**
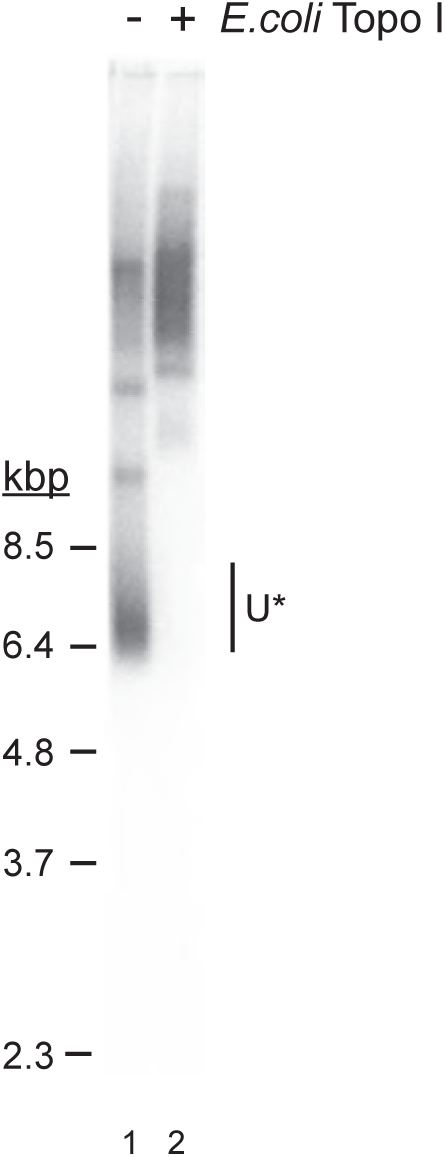
U* is sensitive to relaxation by *E. coli* Topo I. DNA isolated from the reaction analyzed in Figure 6C, lane 5, was either mock-treated (lane 1) or treated with *E. coli* Topo I (lane 2) and analyzed by native agarose gel-electrophoresis.

**Figure S10: Related to Figure 6.**
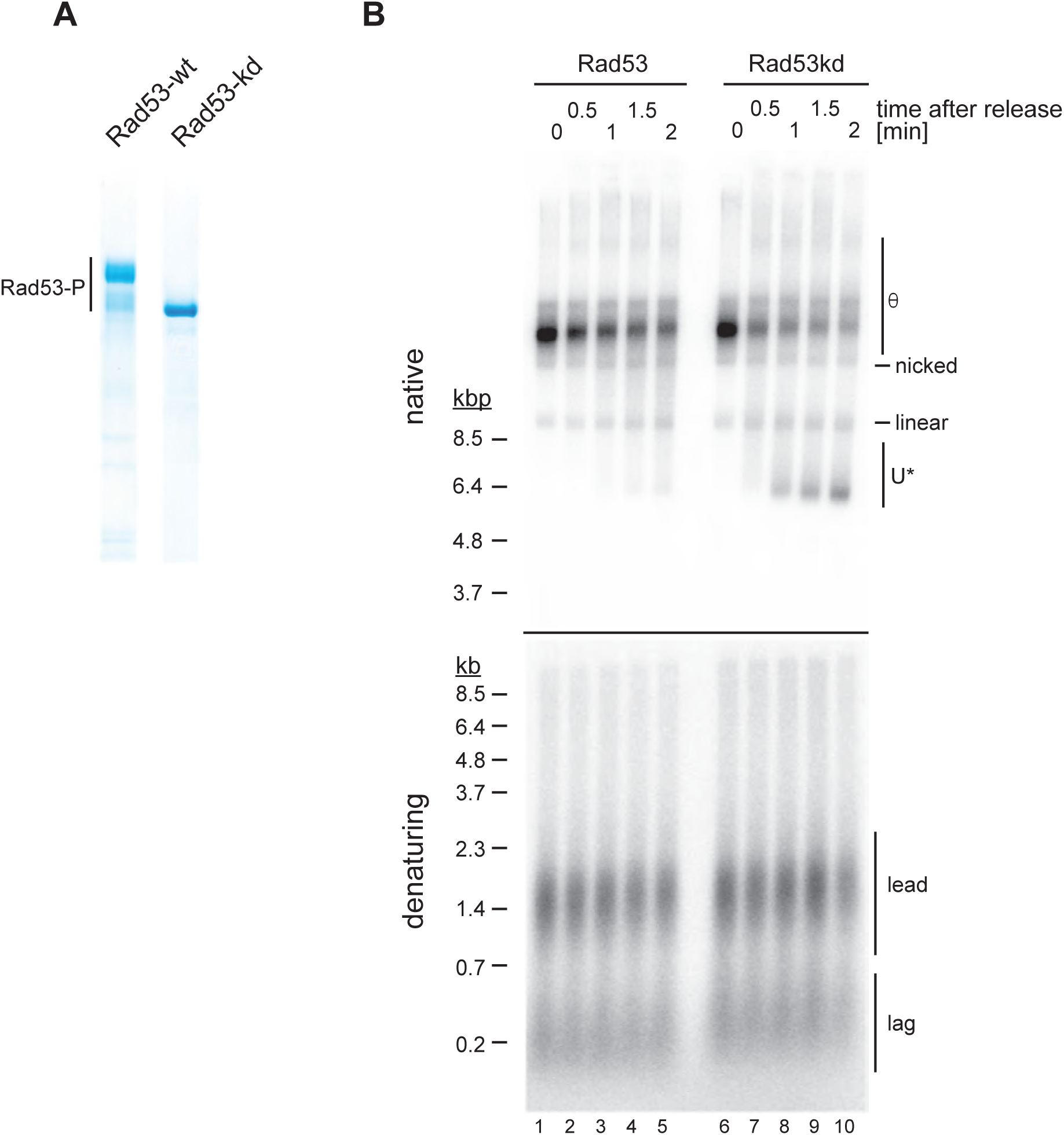
Inhibition of DNA unwinding after CMG uncoupling from DNA synthesis is dependent on the kinase activity of Rad53. (**A**) Purified wild-type and kinase-dead Rad53 (Rad53kd). Rad53-P: Autophosphorylated forms of Rad53. (**B**) Effect of Rad53 kinase activity on U* formation after fork release from topological block in the presence of aphidicolin. Reactions were carried out as in Figure 6C, except that wild-type (lanes 1-5) or kinase-dead (lanes 6-10) Rad53 was added to the reaction prior to fork release.

**Figure S11:**
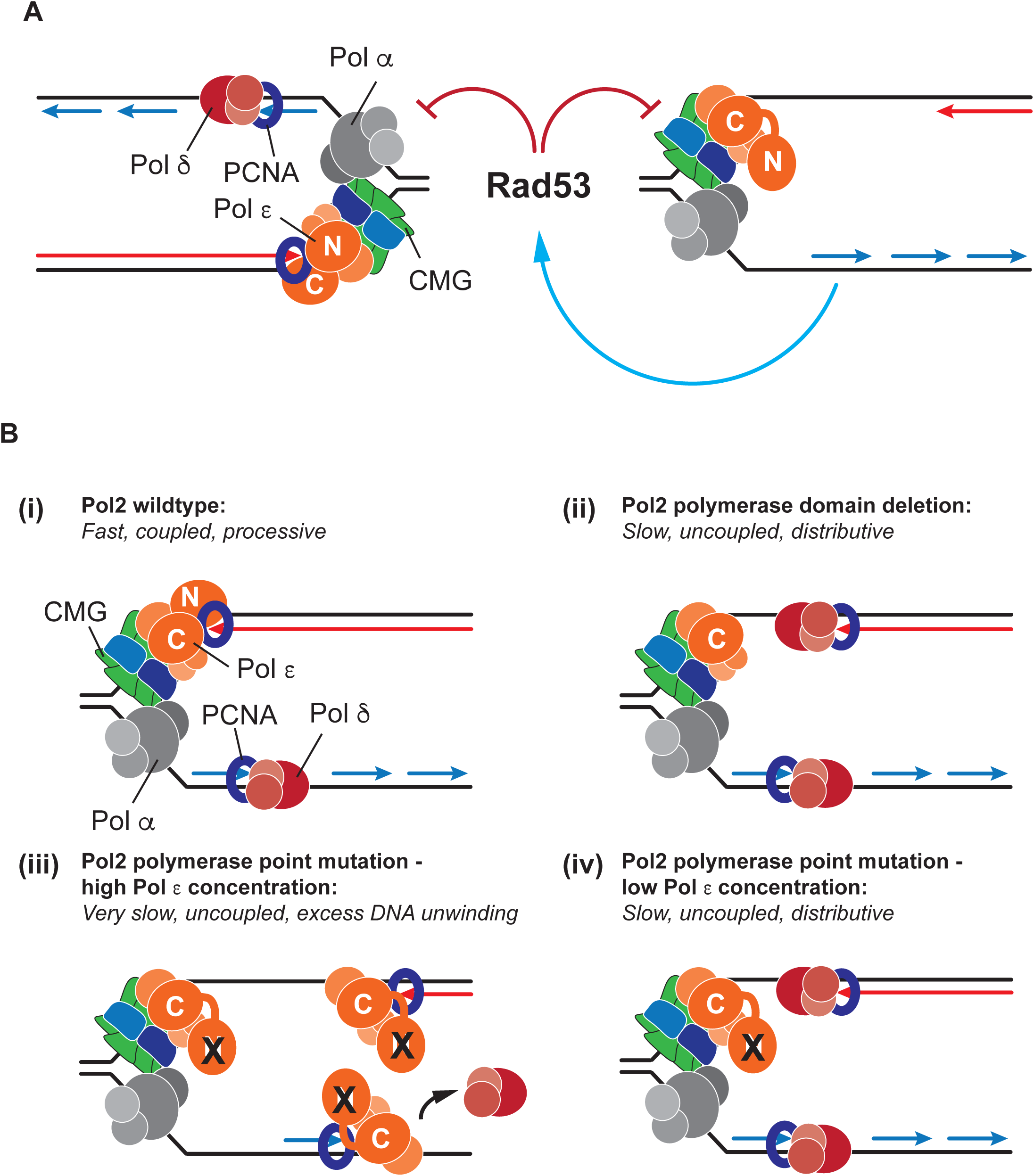
Models for modes of fork progression under conditions of impaired DNA synthesis. (**A**) Cis- and trans-inhibition of replication fork progression upon Rad53 activation. The blue arrow indicates the signaling cascade from RPA/ssDNA to Rad53, red block arrows indicate inhibition of CMG helicase. (**B**) Models for fork progression in the presence of wild-type and catalytically dead Pol ε.

## METHODS

### DNA templates

Plasmids pARS1 (4.8kb) and pARS305 (9.8kb) have been described previously {Devbhandari, 2017 #38;Gros, 2014 #173}. For use in DNA replication reactions, plasmids were purified by sucrose gradient centrifugation following maxiprep column purification (Qiagen). Templates for primer extension reactions were generated by annealing the circular DNA template (M13mp18 ssDNA, NEB) to an oligonucleotide of the sequence 5’-CCCAGTCACGACGTTGTAAAACG-3’. Excess oligonucleotides were removed after the annealing reaction using S-400 spin column chromatography (GE Healthcare).

### Protein purification

Proteins used in this study and a summary of each protein’s purification steps are listed in Table 3. All proteins other than the ones described below were purified as described previously (Devbhandari et al. 2017).

For purifications from yeast, cells carrying galactose-inducible expression constructs were grown at 30°C in YP-GL (YP + 2% glycerol / 2% lactic acid) to a density of 2-4 x 10^7^ cells / ml. Galactose was then added to 2 % and cell growth continued for 4 hours. In the case of Dpb11, cells were arrested in G1 phase for 3 hours by addition of 100 ng / ml alpha factor prior to addition of galactose. Cells were harvested by centrifugation and washed once with 1 M sorbitol / 25 mM Hepes-KOH pH 7.6 followed by a second wash with buffers as indicated. Washed cells were resuspended in 0.5 volumes of respective buffers as indicated and frozen dropwise in liquid nitrogen; the resulting popcorn was stored at −80°C. Frozen popcorn was crushed in a freezer mill (SPEX CertiPrep 6850 Freezer/Mill) for 6 cycles of 2 min at a rate of 15 impacts per second. Extracts were clarified by centrifugation at 42,000 rpm for 45 min (T647.5 rotor). All proteins were stored in aliquots at −80 °C.

#### Csm3·Tof1

Cells were washed with buffer C (25 mM Hepes-KOH pH 7.6 / 0.02 % NP-40 S / 10 % glycerol / 1 mM DTT) / 100 mM NaCl and resuspended in ½ volume of buffer C / 100 mM NaCl / protease inhibitors for preparation of cell popcorn. Cell powder was thawed on ice, one volume of buffer C / 100 mM NaCl was added, and the resulting suspension supplemented with an additional 200 mM NaCl prior to clarification of the extract by ultracentrifugation. The soluble extract was supplemented with 2 mM CaCl_2_ and incubated with calmodulin affinity beads for 3 hours at 4°C. The beads were washed with 10 CVs (column volumes) buffer C / 300 mM NaCl / 2 mM CaCl_2_, and bead-bound protein eluted with 6 CVs buffer C / 300 mM NaCl / 1 mM EDTA / 2 mM EGTA. Eluates were pooled, diluted with an equal volume of buffer C, and fractionated on a Mono Q 5/50 GL column using a gradient of 0.15 – 1 M NaCl over 30 CVs. Peak fractions from the Mono Q step were pooled and fractionated on a 24 ml Superdex 200 gel filtration column equilibrated in buffer C / 300 mM KOAc.

#### Mrc1

Cells were washed with buffer M (25 mM Hepes-KOH pH 7.6 / 1 mM EGTA/ 1 mM EDTA/ 0.02% NP-40 S / 10% glycerol / 0.5 mM DTT) / 100 mM NaCl and resuspended in ½ volumes of buffer M / 100 mM NaCl / protease inhibitors for preparation of popcorn. Cell powder was thawed on ice, one volume of buffer M / 100 mM NaCl was added, and the resulting suspension was supplemented with an additional 300 mM NaCl prior to clarification of the extract by ultracentrifugation. The clarified extract was incubated with M2-agarose anti-FLAG beads at 4°C for 3 hours. The beads were washed with 10 CVs of buffer M / 400 mM NaCl, resuspended in buffer M / 400 mM NaCl / 10 mM Mg-Acetate / 1 mM ATP, and incubated for 10 mins at 4°C, washed again with 10 CV of buffer M / 400 mM NaCl, and bound protein eluted with 1 CV of buffer M / 400 mM NaCl / FLAG peptide (0.5 mg /ml) followed by 2 CVs of buffer M / 400 mM NaCl / FLAG peptide (0.25 mg /ml). Eluates were pooled and diluted with equal volume of buffer M and fractionated on Mono Q column with a salt gradient of 0.2 – 1 M NaCl over 10 CVs. Peak fractions from the Mono Q step were pooled and dialyzed against 25 mM Hepes-KOH pH 7.6 / 1 mM EDTA / 0.02 % NP40-S / 40 % glycerol/ 150 mM NaCl/ 0.5 mM DTT.

#### Dpb11

Cells were washed with buffer D (45 mM Hepes-KOH pH 7.6 / 0.02 % NP-40 S / 10 % glycerol / 1 mM DTT) / 100 mM NaCl, and resuspended in ½ volumes of buffer D/ 100 mM NaCl / protease inhibitors for popcorn preparation. Cell powder was thawed on ice, one volume of buffer D/ 100 mM NaCl was added, the resulting suspension supplemented with an additional 200 mM NaCl and 0.45 % polymin P pH 7.3, and the extract clarified by ultracentrifugation. The clarified extract was supplemented with 2 mM CaCl_2_ and incubated with calmodulin affinity beads for 6 hours at 4°C. The beads were washed with 10 CVs of buffer D / 300 mM NaCl / 2 mM CaCl_2_, and the bead bound protein eluted with 10 CVs buffer D / 300 mM NaCl / 1 mM EDTA/ 2 mM EGTA. Peak fractions were pooled, dialyzed against buffer D / 150 mM NaCl, and fractionated on a Mono Q 5/50 GL column with a salt gradient of 0.15 – 1 M NaCl over 20 CVs. Peak fractions from the Mono Q step were pooled, spin-concentrated, and fractionated on a 24 ml Superdex 200 gel filtration column equilibrated in buffer D / 300 mM KOAc to collect the peak fractions.

#### Sld2

Cells were washed with buffer S (25 mM Hepes-KOH pH 7.6 / 0.02 % NP-40 S / 1 mM EDTA/ 1 mM DTT) / 10 % glycerol / 100 mM KCl and resuspended in ½ volume of buffer S / 10% glycerol/ 100 mM KCl / protease inhibitors for popcorn preparation. Crushed cell powder was thawed on ice, one volume of buffer S / 100 mM KCl was added, and the resulting suspension was supplemented with 400 mM KCl prior to clarification of the extract by ultracentrifugation. The clarified extract was incubated with M2-agarose anti-FLAG beads for 1 hour at 4°C. The beads were washed with 10 CVs of buffer S / 500 mM KCl and bead bound protein was eluted with 1 CV of buffer S / 10 % glycerol 500 mM KCl / FLAG peptide (0.5 mg /ml) followed by 2 CV of buffer S / 10% glycerol / 500 mM KCl / FLAG peptide (0.25 mg / ml). Peak fractions were pooled, dialyzed against buffer S / 40 % glycerol / 350 mM KCl and stored in aliquots.

#### Pol ε variants

Pol ε^WT^ was purified as described previously (Devbhandari et al. 2017), but from asynchronous cells. Pol ε variants (Pol ε^D640A^, Pol ε^DD875,877AA^, Pol ε^Δcat^, Pol ε^D640A-ΔPIP^, Pol ε^exo-^, Pol ε^D640A-exo-^) were purified identically to Pol ε^WT^.

#### Fen1

BL21-Codonplus DE3 (RIL) competent cells (Agilent technologies) were transformed with corresponding vector (Table 1) for Fen1 purification and a culture of transformed cells was grown at 37°C to OD_600_ = 0.6. Cells were then cooled on ice for 30 minutes and expression was induced at 20°C overnight by adding IPTG to 1mM. Cells were harvested via centrifugation and washed twice with autoclaved H_2_0 followed by wash with Buffer F (50 mM Tris-HCl pH 7.5, 1 mM EDTA, 10 % glycerol) / 500 mM NaCl. Then the cells were resuspended in Buffer F / 500 mM NaCl / protease Inhibitor / 1 mM DTT / 0.2 mg per ml lysozyme and lysed by sonication. Cell extract was clarified by centrifugation for 30 minutes at 40,000 rpm in rotor T674.5 (Thermo Fischer Scientific). Clarified extract was incubated for 2 hours with Ni^++^-NTA agarose resin at 4°C. The Ni^++^ -resin was collected by centrifugation in a clinical centrifuge and washed with 10 CVs of Buffer F / 500 mM NaCl / 1 mM DTT followed by wash with 10 CVs of Buffer F / 100 mM NaCl. The bound protein was eluted with 5 CVs of Buffer F / 100 mM NaCl / 1 mM DTT / 100 mM imidazole and incubated with thrombin overnight at 4°C to remove the His-tag. The sample was passed over a Mono Q 5/50 GL column equilibrated in Buffer F / 100 mM NaCl / 1 mM DTT. The flow-through from the Mono Q column, which contains Fen1, was further fractionated by gel-filtration on a 24 ml Superdex 200 column equilibrated in Buffer F / 300 mM K-Acetate / 1 mM DTT. Peak fractions were pooled and stored in aliquots.

#### Rad53

BL21(DE3) pLysS competent cells were transformed with corresponding vector (Table 1) for Rad53 purification and a culture of transformed cells was grown to OD_600_ = 0.6 at 37°C. Cells were then cooled on ice for 15 minutes and expression was induced by adding IPTG to final concentration of 1 mM for 4 hours at 30°C. Cells were harvested by centrifugation and stored at −80°C until further processing. Cells were resuspended in Buffer R (25 mM Tris-HCl pH 7.5, 0.02 % NP-40S, 10 % glycerol) / 1 mM DTT/ 500 mM NaCl / protease inhibitor / 0.1 mg per ml lysozyme and lysed by sonication. Cell debris was removed by centrifugation for 15 minutes in a SS34 rotor at 14,000 rpm, and the resulting clarified extract supplemented with imidazole to final concentration of 10 mM. The extract was incubated for 2 hours with Ni^++^-NTA agarose resin at 4°C. The Ni^++^ - resin was collected by centrifugation and washed with 10 CVs of Buffer R / 10 mM imidazole / 500 mM NaCl / 1 mM DTT. Bound protein was eluted with 5 CVs of Buffer R/ 200 mM imidazole / 500 mM NaCl / 1 mM DTT. Peak fractions were pooled and diluted with 4 volume Buffer R / 1 mM EDTA / 1 mM EGTA / 1 mM DTT to reduce the salt concentration to 100 mM NaCl. The sample was fractionated over a MonoQ 5/50 GL column using a salt gradient of 0.1 - 1 M NaCl in buffer R/ 1 mM EDTA / 1 mM EGTA / 1 mM DTT over 30 CVs. Peak fractions were pooled and further fractionated by gel-filtration on a 24 ml Superdex 200 column equilibrated with 25 mM Hepes-KOH pH 7.6 / 1 mM EDTA / 1 mM EGTA / 10% glycerol / 300 mM K-Acetate / 1 mM DTT. Peak fractions were pooled and stored in aliquots.

#### Rad53 D339A

Rad53 D339A (Rad53kd) was purified identically to Rad53 protein as described above up until the Mono Q 5/50 GL column step. The flow-through from the Mono Q fractionation, which contains the Rad53 D339A, was applied to a Mono S 5/50 GL column and fractionated using a salt gradient of 0.1-1 M NaCl in buffer R/ 1 mM EDTA / 1 mM EGTA / 1 mM DTT over 30 CVs. Mono S peak fractions were pooled and fractionated on a 24 ml Superdex 200 gel filtration column equilibrated with 25 mM Hepes-KOH pH 7.6 / 1 mM EDTA / 1 mM EGTA / 10% glycerol / 300 mM K-Acetate / 1 mM DTT. Peak fractions were pooled and stored in aliquots.

### Standard DNA Replication assay

All steps were carried out at 30°C. First, Mcm2-7 loading was performed in a 10 μl reaction by incubating 10 nM plasmid DNA template, 50 nM ORC, 50 nM Cdc6 and 100 nM Cdt1·Mcm2-7 in a buffer consisting of 25 mM HEPES-KOH (pH7.6), 0.02 % NP-40, 10 mM magnesium acetate, 5 % glycerol, 100 mM potassium acetate, 2 mM DTT, and 5 mM ATP for 15min. DDK was then added to 125 nM and incubation continued for a further 15 min. Unless otherwise stated, DNA replication was initiated by the addition of 15 μl master mix of replication proteins resulting in final concentration of 60 nM Sld3·7, 80 nM Cdc45, 50 nM Clb5·Cdk1, 80 nM GINS, 30 nM Dpb11, 30nM Pol ε, 80 nM Sld2, 120 nM RPA, 60 nM Pol α, 35 nM Ctf4, 20 nM RFC, 70 nM PCNA, 4 nM Pol δ, 20 nM Csm3·Tof1, 20 nM Mrc1, 15 nM Mcm10, 30 nM Top1, and 30 nM Top2. The final replication reaction also included 0.2 mg / ml BSA, 40 μM each dATP / dGTP / dTTP / dCTP, 200 μM each GTP / CTP /UTP, 66 nM α^32^P-dATP (3,000 Ci / mmol), 16 mM creatine phosphate, 0.04 mg / ml creatine kinase, 190 nM potassium acetate, 20 mM sodium chloride, and 15 mM potassium chloride. Reactions were terminated at times indicated in the respective figures by adding 40 mM EDTA, 1.6 U Proteinase K, and 0.25 % SDS, and incubating the mix at 37°C for 30 minutes. DNA was isolated by phenol / chloroform extraction and unincorporated nucleotides removed with G-50 spin columns equilibrated in TE buffer. The sample was then fractionated on a 0.8% alkaline agarose gel (30 mM NaOH and 2 mM EDTA) or 0.8% native agarose gel (in TAE) dried, and imaged using Typhoon FLA 7000. Quantification of the gel images was performed using the ImageJ software.

### Modified replication assays

In Figure 1A, the master mix of replication proteins included Fen1 and Cdc9 as indicated, resulting in final concentrations of 15 nM and 12 nM, respectively. In Fig 1E, the concentration of Pol ε or its variants was 15 nM. For the dNTP titration experiment in Fig 5A (lanes 1-3) the final concentration of α^32^P-dATP was 3.3 nM; the shown experiment was performed in the absence of Csm3·Tof1 and Mrc1. For reactions with limiting dNTPs in Fig 5B and Fig S8, the final concentration of dNTPs was 250 nM each.

To stall replisomes in absence of topoisomerase (Figures 6 and 7), replication was initiated with master mix devoid of Top1 and Top2 and continued for 10 minutes, followed by simultaneous addition of Top1, Top2 and aphidicolin (or DMSO as control), as indicated, to final concentrations of 30 nM, 30 nM and 30 μM respectively. In experiments involving Rad53, 7 minutes after the addition of master mix (lacking Top1 and Top2), Rad53 was added to final concentration of 100 nM and incubation was continued for an additional 7 minutes prior to addition of either Top1 and Top2 with or without aphidicolin.

For protein titration experiments, the stock of the corresponding protein was serially diluted in respective storage buffer and equal volumes of the protein dilutions were added to the initiation master mix. For time course assays, equal volumes of a replication reaction were removed from a single master reaction and terminated at indicated times as described above for standard replication assay.

### dNTP pulse-chase replication reactions

Reactions in Figure 2 were carried out identically to the standard replication assay with the exception that the dATP concentration was lowered to 2 μM during the pulse and the addition of cold dATP to final concentration of 1 mM for the chase. Reactions were pulse-labeled for 10 mins (Figure 2B) or 2.5 mins (Figure 2D). In Figures 5B and 5C, the final concentration of cold dATP was raised to 500 μM after 5 minutes.

### Primer extension assay

Primer extension reactions were performed at 30°C in polymerization buffer (25 mM Tris-HCl pH 7.5, 8 mM Mg-Acetate, 50 mM K-glutamate, 5 % glycerol). First, 1 nM of DNA template was incubated with 1 mM ATP, 1 mM DTT, 80 μM dATP, 80 μM dGTP, 80 μM dCTP and 400 nM of RPA for 5 mins. PCNA and RFC were then added to 70 nM and 4 nM respectively and incubation was continued for 5 more minutes. Then, 4 nM Pol δ and 33 nM α^32^P-dATP (3,000 Ci / mmol) was added to the reaction resulting in a primer extension by 9 bp (due to lack of dTTP). After 5 mins, variants of Pol ε (to final concentrations as indicated) and 80 μM dTTP were added to the mix and the extension was continued for 3 minutes. The reactions were stopped by the addition of EDTA and SDS to final concentrations of 40 mM and 0.25 %, respectively. Products were fractionated on a 0.8 % alkaline agarose gel (30 mM NaOH and 2 mM EDTA), dried, and imaged using Typhoon FLA 7000. Quantification of the gel images was performed using the ImageJ software.

### CMG helicase assay

All steps were carried out at 30°C. First, Mcm2-7 loading was performed by incubating 140 fmol of plasmid DNA template with 50 nM ORC, 50 nM Cdc6 and 100 nM Cdt1·Mcm2-7 in a buffer consisting of 25 mM HEPES-KOH (pH7.6), 0.02 % NP-40, 10 mM magnesium acetate, 5 % glycerol, 100 mM potassium acetate, 2 mM DTT, 5 mM ATP, 40 mM creatine phosphate and 0.1 mg / ml creatine kinase for 15 minutes. DDK was then added to 125 nM and incubation continued for a further 15 minutes. Then a master mix of proteins for CMG activation was added to the reaction resulting in final concentration of 75 nM Sld3·7, 100 nM Cdc45, 60 nM CDK, 100 nM GINS, 40 nM Dpb11, 30 nM Pol ε, 100 nM Sld2, 135 nM RPA, 15 nM Mcm10, and 0.4 mg / ml BSA. The reaction mix was incubated for 10 minutes and Rad53 was then added, as indicated, to a final concentration of 100 nM. Incubation was continued for 5 minutes and Top1 was added to final concentration of 30 nM. Aliquots of the reaction were removed at indicated times and terminated by adding 25 mM EDTA, 1.6 U Proteinase K and 0.25% SDS, and incubating the mix at 42°C for 20 minutes. DNA was isolated by phenol / chloroform extraction and unincorporated nucleotides removed by centrifugation through G-50 spin columns equilibrated with TE buffer. Products were fractionated on a 1.0 % native agarose gel (in TAE) and visualized by staining the gel with 1x TAE consisting of 0.5 μg / ml of ethidium bromide for 1 hour followed by de-staining with 1x TAE/ 1 mM magnesium sulfate for 1 hour.

### Plasmid Shuffle assay with Pol ε variants

Yeast strain T-1925-5 (W303 *leu2-3,112 trp1-1 can1-100 ura3-1 ade2-1 his3-11,15 pol2*Δ*::KAN pRS416-Pol2-3HA::*), a kind gift of Xiaolan Zhao (MSKCC), was transformed with respective pRS315G-POL2 tester plasmids and grown on selective minimal plates lacking uracil and leucine. Resulting transformants were re-streaked on minimal plates lacking leucine and grown for 24 hours at 30°C to allow loss of pRS416-POL2^wt^. Transformants were then restreaked on minimal plates lacking leucine and containing 0.1 % 5-fluoroorotic acid (5-FOA), 2 % raffinose, and either 0.05% or 1% galactose, as indicated. For the Western blot analysis in Figure 4A, transformants were grown to the density of 2 x 10^7^ cells / ml in selective media lacking leucine and uracil and containing 2 % raffinose. Galactose was then added to final concentration of either 0.05 % or 1 %, and cells collected after 3 hours and lysed and processed for western blotting.

### Quantification and statistical analysis

Data were quantified and statistically analyzed using Image J and Graphpad Prism 7.0 softwares, respectively. For the pulse-chase experiments in Figure 2, lane profiles of the gels were generated using the Image J and the maximum product length was appropriated as the intersection of two straight lines, corresponding to the straight line manually fitted to lane background and second the straight line manually fitted to the leading edge of the leading strand population. A standard curve generated from molecular marker was used to convert the point of intersection to length in base pairs. Replication rates were calculated by fitting the data into a linear regression and calculating the slope of the regression.

For the quantification of relative DNA synthesis in Figure 3B, total DNA synthesis per lane was first calculated using the lane profiles and divided by the amount of DNA synthesis in lane 1 (without Pol ε). In Figure S2A, the percent dissolution was calculated as the fraction of full length linear product relative to total replication products in each lane.

### Plasmids used in this study (Table 1)

| Name | Insert |
| --- | --- |
| pRS306G-CSM3-CBP/TOF1 | CSM3 (amplified from *S. cerevisiae* W303, C-terminal CBP tag), TOF1 (amplified from *S. cerevisiae* W303) |
| pRS305G-MRC1-FLAG | MRC1 (synthetic ORF, C-terminal 3x FLAG tag) |
| pRS306G-DPB4/CBP-POL2<sup>DD875,877AA</sup> | DPB4 (synthetic ORF) CBP-POL2 (synthetic ORF with D875A D877A mutations, N-terminal CBP tag) |
| pRS306G-DPB4/CBP-POL2<sup>Δcat</sup> | DPB4 (synthetic ORF) CBP-POL2 (synthetic ORF containing amino acids 1263-2222, N-terminal CBP tag) |
| pRS306G-DPB4/CBP-POL2<sup>D640A-ΔPIP</sup> | DPB4 (synthetic ORF) CBP-POL2 (synthetic ORF with D640A, F1199A, F1200A mutations, N-terminal CBP tag) |
| pRS306G-DPB4/CBP-POL2<sup>exo-</sup> | DPB4 (synthetic ORF) CBP-POL2 (synthetic ORF with D290A E292A mutations, N-terminal CBP tag) |
| pRS306G-DPB4/CBP-POL2<sup>D640A-exo-</sup> | DPB4 (synthetic ORF) CBP-POL2 (synthetic ORF with D640A, D290A, E292A mutations N-terminal CBP tag) |
| pRS315G-CBP-POL2 | GAL1,10 promoter + CBP-POL2 (synthetic ORF, N-terminal CBP tag) + PGK1 3' UTR inserted into pRS315 |
| pRS315G-CBP-POL2-16 | GAL1,10 promoter + CBP-*pol2-16* (synthetic ORF, N-terminal CBP tag) + PGK1 3' UTR inserted into pRS315 |
| pRS315G-CBP-POL2<sup>DD875,877AA</sup> | GAL1,10 promoter + CBP-*pol2-DD875,877AA* (synthetic ORF, N-terminal CBP tag) + PGK1 3' UTR inserted into pRS315 |
| pRS315G-CBP-POL2<sup>Δcat</sup> | GAL1,10 promoter + CBP-*pol2Δcat* (synthetic ORF, N-terminal CBP tag) + PGK1 3' UTR inserted into pRS315 |
| pRS315G-CBP-POL2<sup>D640A</sup> | GAL1,10 promoter + CBP-*pol2-D640A* (synthetic ORF, N-terminal CBP tag) + PGK1 3' UTR inserted into pRS315 |
| pRS315G-CBP-POL2<sup>D640A-ΔPIP</sup> | GAL1,10 promoter + CBP-*pol2-D640A-FF1199,1200AA* (synthetic ORF, N-terminal CBP tag) + PGK1 3' UTR inserted into pRS315 |
| pRS315G-CBP-POL2<sup>DD875,877AA-ΔPIP</sup> | GAL1,10 promoter + CBP-*pol2-DD875,877AA-FF1199,1200AA* (synthetic ORF, N-terminal CBP tag) + PGK1 3' UTR inserted into pRS315 |
| pRS315G-CBP-POL2<sup>D640A-exo-</sup> | GAL1,10 promoter + CBP-*pol2-D640A-D290A-E292A* (synthetic ORF, N-terminal CBP tag) + PGK1 3' UTR inserted into pRS315 |
| pET28a-FEN1-6xHis | FEN1 (amplified from *S. cerevisiae* W303, C-terminal 6x His tag) |
| Pet15b-6xHis-RAD53 | RAD53 (amplified from *S. cerevisiae* W303, N-terminal 6x His tag) |
| Pet15b-6xHis-RAD53<sup>D339A</sup> | RAD53 with D339A mutation (amplified from *S. cerevisiae* W303, N-terminal 6x His tag) |

### Yeast strains used in this study (Table 2)

| Name | Genotype |
| --- | --- |
| **YDR137** | *MATa ade2-1 ura3-1 his3-11,15 trp1-1 leu2-3,112 can1-100*<br>*pep4::kanMX bar::hphNAT1 Gal-GAL4 (HIS3) Gal-CSM3-CBP / TOF1 (URA3)* |
| **YDR163** | *MATa ade2-1 ura3-1 his3-11,15 trp1-1 leu2-3,112 can1-100*<br>*pep4::kanMX bar::hphNAT1 Gal-GAL4 (HIS3) Gal-MRC1-FLAG (LEU2)* |
| **YDR110** | *MATa ade2-1 ura3-1 his3-11,15 trp1-1 leu2-3,112 can1-100*<br>*pep4::kanMX bar::hphNAT1 Gal-GAL4 (HIS3) Gal-DPB11-CBP (URA3)* |
| **YSD13** | *MATa ade2-1 ura3-1 his3-11,15 trp1-1 leu2-3,112 can1-100*<br>*pep4::kanMX bar::hphNAT1 Gal-GAL4 (HIS3) Gal-SLD2-FLAG (URA3)* |
| **YDR166** | *MATa ade2-1 ura3-1 his3-11,15 trp1-1 leu2-3,112 can1-100*<br>*pep4::kanMX bar::hphNAT1 Gal-GAL4 (HIS3) Gal-DPB2/DPB3 (LEU2)*<br>*Gal-CBP-pol2-D875A, D877A / DPB4 (URA3)* |
| **YDR172** | *MATa ade2-1 ura3-1 his3-11,15 trp1-1 leu2-3,112 can1-100*<br>*pep4::kanMX bar::hphNAT1 Gal-GAL4 (HIS3) Gal-DPB2 / DPB3 (LEU2)*<br>*Gal-CBP-pol2-D640A-F1199A,F1200A / DPB4 (URA3)* |
| **YDR165** | *MATa ade2-1 ura3-1 his3-11,15 trp1-1 leu2-3,112 can1-100*<br>*pep4::kanMX bar::hphNAT1 Gal-GAL4 (HIS3) Gal-DPB2 / DPB3 (LEU2)*<br>*Gal-CBP-pol2-Δ1-1262 / DPB4 (URA3)* |
| **YSD20** | *MATa ade2-1 ura3-1 his3-11,15 trp1-1 leu2-3,112 can1-100*<br>*pep4::kanMX bar::hphNAT1 Gal-GAL4 (HIS3) Gal-DPB2 / DPB3 (LEU2)*<br>*Gal-CBP-pol2-D640A-D290A-E292A / DPB4 (URA3)* |
| **YDR144** | *MATa ade2-1 ura3-1 his3-11,15 trp1-1 leu2-3,112 can1-100*<br>*pep4::kanMX bar::hphNAT1 Gal-GAL4 (HIS3) Gal-DPB2 / DPB3 (LEU2)*<br>*Gal-CBP-pol2 -D290A,E292A / DPB4 (URA3)* |
| **T1925-5** | *W303 leu2-3,112 trp1-1 can1-100 ura3-1 ade2-1 his3-11,15*<br>*pol2Δ::KAN pRS416-Pol2-3HA::KAN* |

### Summary of purification steps for individual proteins (Table 3)

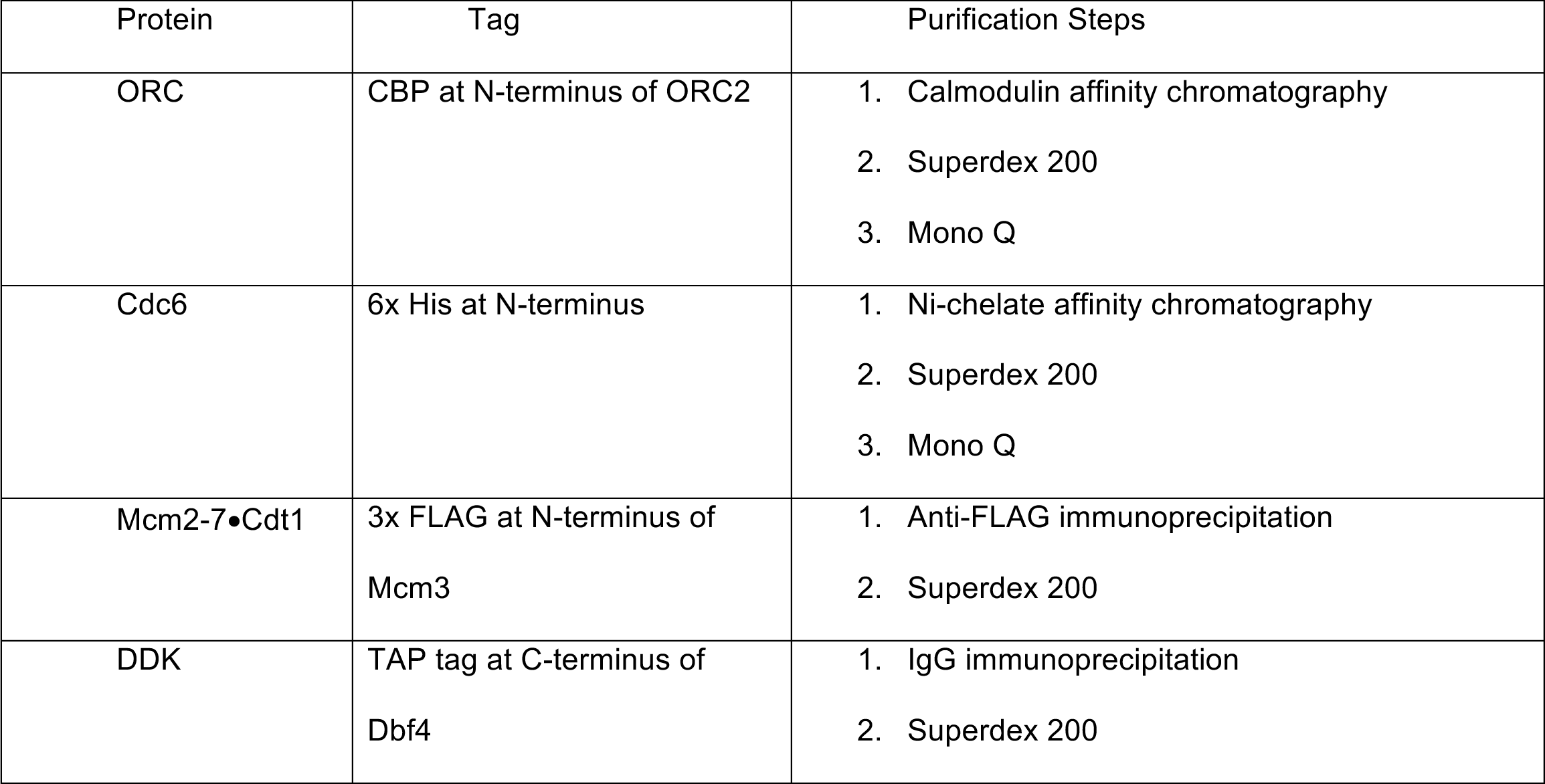
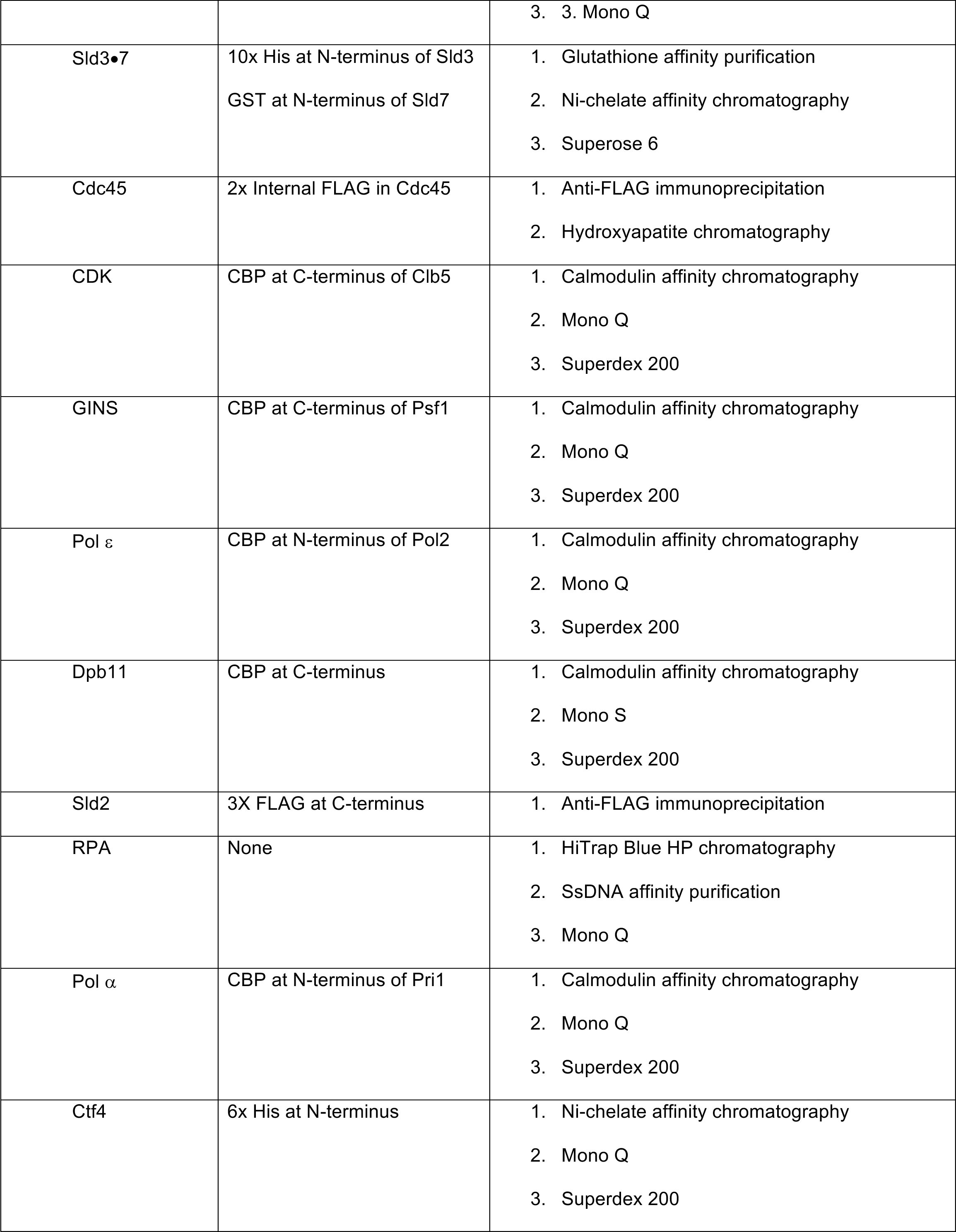
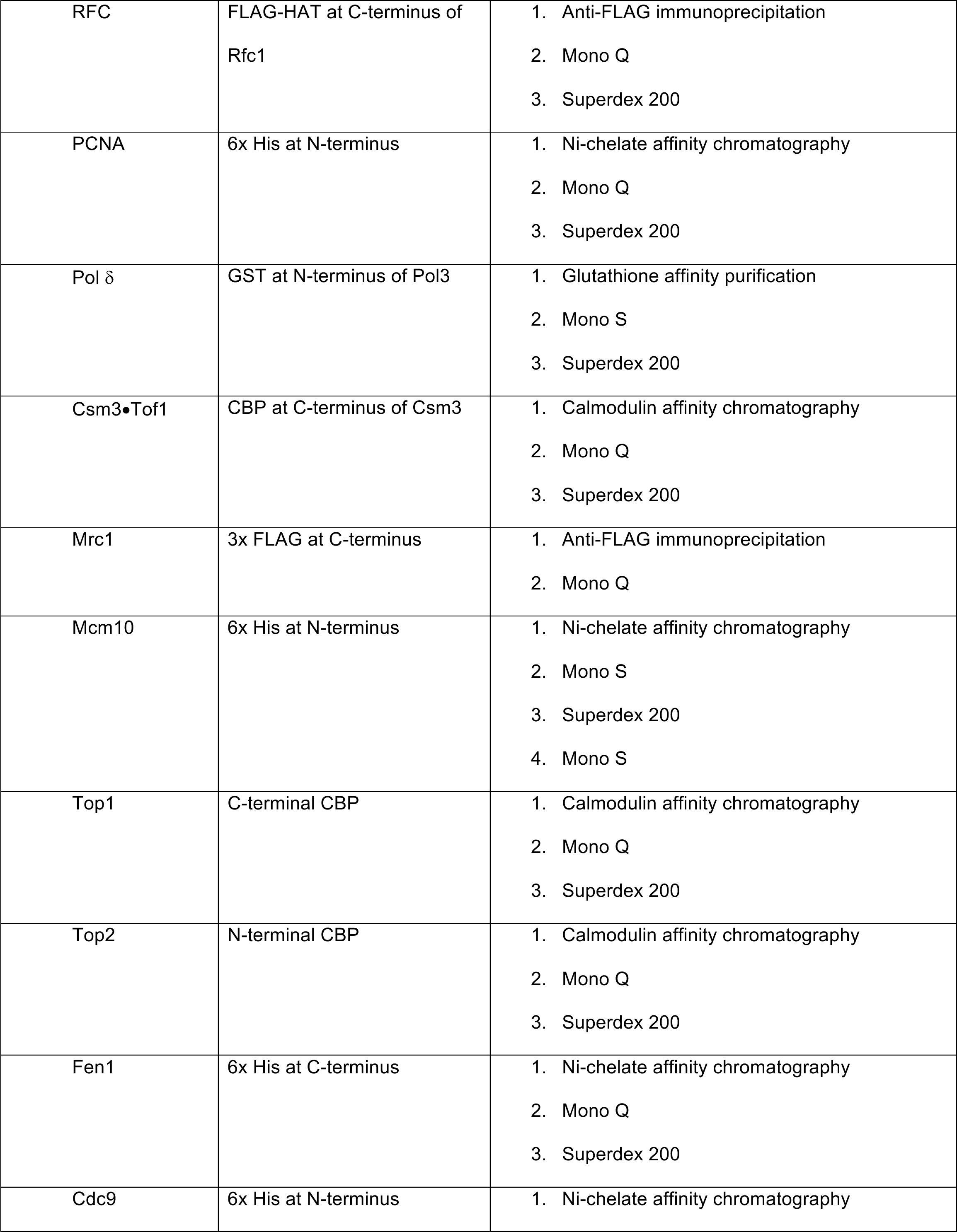
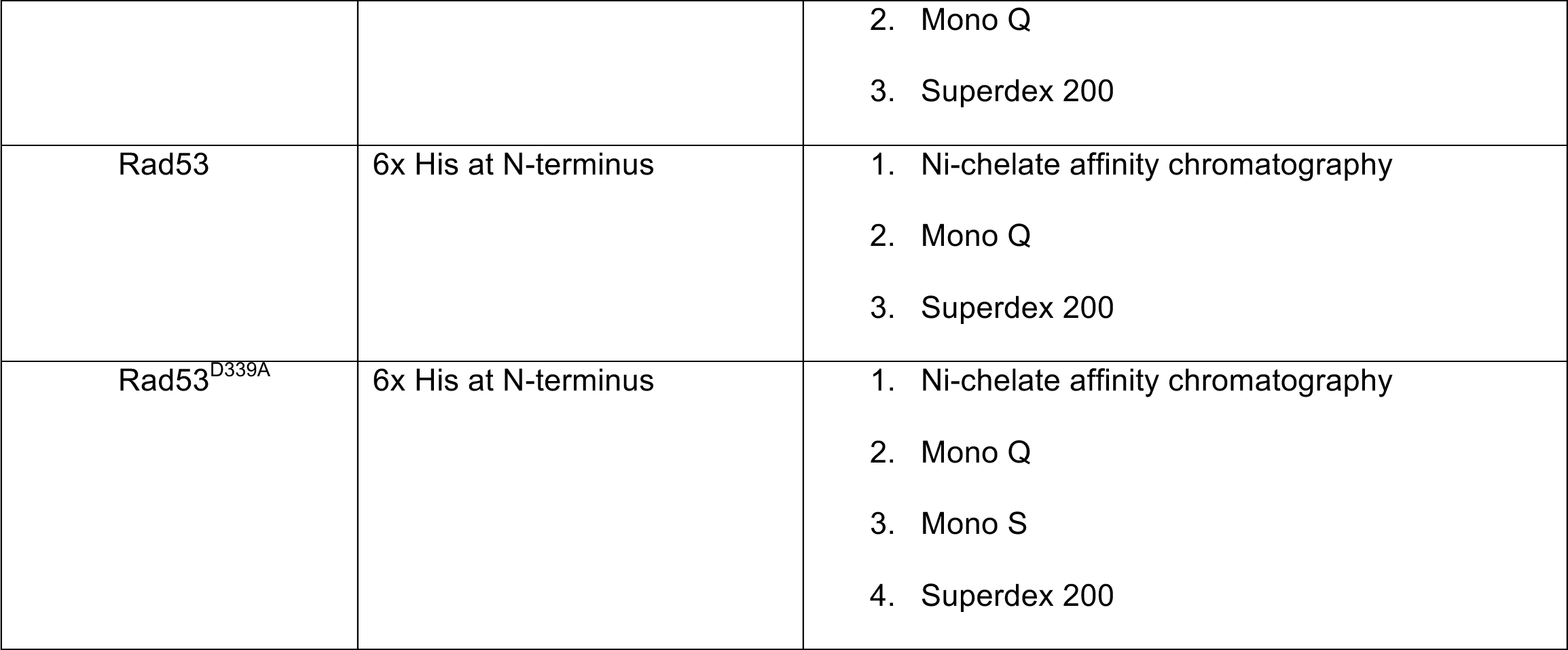

## REFERENCES

1. Bell, S.P. & Labib, K. Chromosome Duplication in Saccharomyces cerevisiae. Genetics 203, 1027–67 (2016).

2. Burgers, P.M.J. & Kunkel, T.A. Eukaryotic DNA Replication Fork. Annu Rev Biochem 86, 417–438 (2017).

3. Lujan, S.A., Williams, J.S. & Kunkel, T.A. DNA Polymerases Divide the Labor of Genome Replication. Trends Cell Biol 26, 640–654 (2016).

4. Li, H. & O’Donnell, M.E. The Eukaryotic CMG Helicase at the Replication Fork: Emerging Architecture Reveals an Unexpected Mechanism. Bioessays 40(2018).

5. Tahirov, T.H., Makarova, K.S., Rogozin, I.B., Pavlov, Y.I. & Koonin, E.V. Evolution of DNA polymerases: an inactivated polymerase-exonuclease module in Pol epsilon and a chimeric origin of eukaryotic polymerases from two classes of archaeal ancestors. Biol Direct 4, 11 (2009).

6. Goswami, P. et al. Structure of DNA-CMG-Pol epsilon elucidates the roles of the non-catalytic polymerase modules in the eukaryotic replisome. Nat Commun 9, 5061 (2018).

7. Hogg, M. et al. Structural basis for processive DNA synthesis by yeast DNA polymerase varepsilon. Nat Struct Mol Biol 21, 49–55 (2014).

8. Dua, R., Levy, D.L. & Campbell, J.L. Analysis of the essential functions of the C-terminal protein/protein interaction domain of Saccharomyces cerevisiae pol epsilon and its unexpected ability to support growth in the absence of the DNA polymerase domain. J Biol Chem 274, 22283–8 (1999).

9. Kesti, T., Flick, K., Keranen, S., Syvaoja, J.E. & Wittenberg, C. DNA polymerase epsilon catalytic domains are dispensable for DNA replication, DNA repair, and cell viability. Mol Cell 3, 679–85 (1999).

10. Garbacz, M.A. et al. Evidence that DNA polymerase delta contributes to initiating leading strand DNA replication in Saccharomyces cerevisiae. Nat Commun 9, 858 (2018).

11. Garbacz, M.A. et al. The absence of the catalytic domains of Saccharomyces cerevisiae DNA polymerase strongly reduces DNA replication fidelity. Nucleic Acids Res 47, 3986–3995 (2019).

12. Georgescu, R.E. et al. Mechanism of asymmetric polymerase assembly at the eukaryotic replication fork. Nat Struct Mol Biol 21, 664–70 (2014).

13. Yeeles, J.T.P., Janska, A., Early, A. & Diffley, J.F.X. How the Eukaryotic Replisome Achieves Rapid and Efficient DNA Replication. Mol Cell 65, 105–116 (2017).

14. Schauer, G.D. & O’Donnell, M.E. Quality control mechanisms exclude incorrect polymerases from the eukaryotic replication fork. Proc Natl Acad Sci U S A 114, 675–680 (2017).

15. Delagoutte, E. & von Hippel, P.H. Molecular mechanisms of the functional coupling of the helicase (gp41) and polymerase (gp43) of bacteriophage T4 within the DNA replication fork. Biochemistry 40, 4459–77 (2001).

16. Kim, S., Dallmann, H.G., McHenry, C.S. & Marians, K.J. Coupling of a replicative polymerase and helicase: a tau-DnaB interaction mediates rapid replication fork movement. Cell 84, 643–50 (1996).

17. Manosas, M., Spiering, M.M., Ding, F., Croquette, V. & Benkovic, S.J. Collaborative coupling between polymerase and helicase for leading-strand synthesis. Nucleic Acids Res 40, 6187–98 (2012).

18. Stano, N.M. et al. DNA synthesis provides the driving force to accelerate DNA unwinding by a helicase. Nature 435, 370–3 (2005).

19. Burnham, D.R., Kose, H.B., Hoyle, R.B. & Yardimci, H. The mechanism of DNA unwinding by the eukaryotic replicative helicase. Nat Commun 10, 2159 (2019).

20. Sparks, J.L. et al. The CMG Helicase Bypasses DNA-Protein Cross-Links to Facilitate Their Repair. Cell 176, 167–181 e21 (2019).

21. Taylor, M.R.G. & Yeeles, J.T.P. Dynamics of Replication Fork Progression Following Helicase-Polymerase Uncoupling in Eukaryotes. J Mol Biol 431, 2040–2049 (2019).

22. Taylor, M.R.G. & Yeeles, J.T.P. The Initial Response of a Eukaryotic Replisome to DNA Damage. Mol Cell 70, 1067–1080 e12 (2018).

23. Walter, J. & Newport, J. Initiation of eukaryotic DNA replication: origin unwinding and sequential chromatin association of Cdc45, RPA, and DNA polymerase alpha. Mol Cell 5, 617–27 (2000).

24. Gan, H. et al. Checkpoint Kinase Rad53 Couples Leading- and Lagging-Strand DNA Synthesis under Replication Stress. Mol Cell 68, 446–455 e3 (2017).

25. Katou, Y. et al. S-phase checkpoint proteins Tof1 and Mrc1 form a stable replication-pausing complex. Nature 424, 1078–83 (2003).

26. Byun, T.S., Pacek, M., Yee, M.C., Walter, J.C. & Cimprich, K.A. Functional uncoupling of MCM helicase and DNA polymerase activities activates the ATR-dependent checkpoint. Genes Dev 19, 1040–52 (2005).

27. Zou, L. & Elledge, S.J. Sensing DNA damage through ATRIP recognition of RPA-ssDNA complexes. Science 300, 1542–8 (2003).

28. Giannattasio, M. & Branzei, D. S-phase checkpoint regulations that preserve replication and chromosome integrity upon dNTP depletion. Cell Mol Life Sci 74, 2361–2380 (2017).

29. Saldivar, J.C., Cortez, D. & Cimprich, K.A. The essential kinase ATR: ensuring faithful duplication of a challenging genome. Nat Rev Mol Cell Biol 18, 622–636 (2017).

30. Lanz, M.C., Dibitetto, D. & Smolka, M.B. DNA damage kinase signaling: checkpoint and repair at 30 years. EMBO J, e101801 (2019).

31. Branzei, D. & Foiani, M. Maintaining genome stability at the replication fork. Nat Rev Mol Cell Biol 11, 208–19 (2010).

32. De Piccoli, G. et al. Replisome stability at defective DNA replication forks is independent of S phase checkpoint kinases. Mol Cell 45, 696–704 (2012).

33. Dungrawala, H. et al. The Replication Checkpoint Prevents Two Types of Fork Collapse without Regulating Replisome Stability. Mol Cell 59, 998–1010 (2015).

34. Cortez, D. Preventing replication fork collapse to maintain genome integrity. DNA Repair (Amst) 32, 149–157 (2015).

35. Segurado, M. & Diffley, J.F. Separate roles for the DNA damage checkpoint protein kinases in stabilizing DNA replication forks. Genes Dev 22, 1816–27 (2008).

36. Sogo, J.M., Lopes, M. & Foiani, M. Fork reversal and ssDNA accumulation at stalled replication forks owing to checkpoint defects. Science 297, 599–602 (2002).

37. Devbhandari, S., Jiang, J., Kumar, C., Whitehouse, I. & Remus, D. Chromatin Constrains the Initiation and Elongation of DNA Replication. Mol Cell 65, 131–141 (2017).

38. Deegan, T.D., Baxter, J., Ortiz Bazan, M.A., Yeeles, J.T.P. & Labib, K.P.M. Pif1-Family Helicases Support Fork Convergence during DNA Replication Termination in Eukaryotes. Mol Cell 74, 231–244 e9 (2019).

39. Baker, T.A., Sekimizu, K., Funnell, B.E. & Kornberg, A. Extensive unwinding of the plasmid template during staged enzymatic initiation of DNA replication from the origin of the Escherichia coli chromosome. Cell 45, 53–64 (1986).

40. Dean, F.B. et al. Simian virus 40 (SV40) DNA replication: SV40 large T antigen unwinds DNA containing the SV40 origin of replication. Proc Natl Acad Sci U S A 84, 16–20 (1987).

41. Wold, M.S., Li, J.J. & Kelly, T.J. Initiation of simian virus 40 DNA replication in vitro: large-tumor-antigen- and origin-dependent unwinding of the template. Proc Natl Acad Sci U S A 84, 3643–7 (1987).

42. Langston, L.D. et al. CMG helicase and DNA polymerase epsilon form a functional 15-subunit holoenzyme for eukaryotic leading-strand DNA replication. Proc Natl Acad Sci U S A 111, 15390–5 (2014).

43. Sun, J. et al. The architecture of a eukaryotic replisome. Nat Struct Mol Biol 22, 976–82 (2015).

44. Zhou, J.C. et al. CMG-Pol epsilon dynamics suggests a mechanism for the establishment of leading-strand synthesis in the eukaryotic replisome. Proc Natl Acad Sci U S A 114, 4141–4146 (2017).

45. Douglas, M.E., Ali, F.A., Costa, A. & Diffley, J.F.X. The mechanism of eukaryotic CMG helicase activation. Nature 555, 265–268 (2018).

46. Georgescu, R.E. et al. Reconstitution of a eukaryotic replisome reveals suppression mechanisms that define leading/lagging strand operation. Elife 4, e04988 (2015).

47. Chilkova, O. et al. The eukaryotic leading and lagging strand DNA polymerases are loaded onto primer-ends via separate mechanisms but have comparable processivity in the presence of PCNA. Nucleic Acids Res 35, 6588–97 (2007).

48. Dua, R., Levy, D.L., Li, C.M., Snow, P.M. & Campbell, J.L. In vivo reconstitution of Saccharomyces cerevisiae DNA polymerase epsilon in insect cells. Purification and characterization. J Biol Chem 277, 7889–96 (2002).

49. Ganai, R.A. & Johansson, E. DNA Replication-A Matter of Fidelity. Mol Cell 62, 745–55 (2016).

50. Jain, R. et al. Crystal structure of yeast DNA polymerase epsilon catalytic domain. PLoS One 9, e94835 (2014).

51. Ganai, R.A., Bylund, G.O. & Johansson, E. Switching between polymerase and exonuclease sites in DNA polymerase epsilon. Nucleic Acids Res 43, 932–42 (2015).

52. Ganai, R.A., Zhang, X.P., Heyer, W.D. & Johansson, E. Strand displacement synthesis by yeast DNA polymerase epsilon. Nucleic Acids Res 44, 8229–40 (2016).

53. Deegan, T.D., Yeeles, J.T. & Diffley, J.F. Phosphopeptide binding by Sld3 links Dbf4-dependent kinase to MCM replicative helicase activation. EMBO J 35, 961–73 (2016).

54. Lopez-Mosqueda, J. et al. Damage-induced phosphorylation of Sld3 is important to block late origin firing. Nature 467, 479–83 (2010).

55. Zegerman, P. & Diffley, J.F. Checkpoint-dependent inhibition of DNA replication initiation by Sld3 and Dbf4 phosphorylation. Nature 467, 474–8 (2010).

56. Szyjka, S.J., Viggiani, C.J. & Aparicio, O.M. Mrc1 is required for normal progression of replication forks throughout chromatin in S. cerevisiae. Mol Cell 19, 691–7 (2005).

57. Tourriere, H., Versini, G., Cordon-Preciado, V., Alabert, C. & Pasero, P. Mrc1 and Tof1 promote replication fork progression and recovery independently of Rad53. Mol Cell 19, 699–706 (2005).

58. Smolka, M.B., Albuquerque, C.P., Chen, S.H. & Zhou, H. Proteome-wide identification of in vivo targets of DNA damage checkpoint kinases. Proc Natl Acad Sci U S A 104, 10364–9 (2007).

59. Alcasabas, A.A. et al. Mrc1 transduces signals of DNA replication stress to activate Rad53. Nat Cell Biol 3, 958–65 (2001).

60. Graham, J.E., Marians, K.J. & Kowalczykowski, S.C. Independent and Stochastic Action of DNA Polymerases in the Replisome. Cell 169, 1201–1213 e17 (2017).

61. Burgers, P.M. Saccharomyces cerevisiae replication factor C. II. Formation and activity of complexes with the proliferating cell nuclear antigen and with DNA polymerases delta and epsilon. J Biol Chem 266, 22698–706 (1991).

62. Daigaku, Y. et al. A global profile of replicative polymerase usage. Nat Struct Mol Biol 22, 192–198 (2015).

63. Aria, V. & Yeeles, J.T.P. Mechanism of Bidirectional Leading-Strand Synthesis Establishment at Eukaryotic DNA Replication Origins. Mol Cell (2018).

64. Stillman, B. Reconsidering DNA Polymerases at the Replication Fork in Eukaryotes. Mol Cell 59, 139–41 (2015).

65. Sun, B. et al. ATP-induced helicase slippage reveals highly coordinated subunits. Nature 478, 132–5 (2011).

66. Wasserman, M.R., Schauer, G.D., O’Donnell, M.E. & Liu, S. Replication Fork Activation Is Enabled by a Single-Stranded DNA Gate in CMG Helicase. Cell 178, 600–611 e16 (2019).

67. Cotta-Ramusino, C. et al. Exo1 processes stalled replication forks and counteracts fork reversal in checkpoint-defective cells. Mol Cell 17, 153–9 (2005).

68. Toledo, L.I. et al. ATR prohibits replication catastrophe by preventing global exhaustion of RPA. Cell 155, 1088–103 (2013).

69. Neelsen, K.J. & Lopes, M. Replication fork reversal in eukaryotes: from dead end to dynamic response. Nat Rev Mol Cell Biol 16, 207–20 (2015).

70. Can, G., Kauerhof, A.C., Macak, D. & Zegerman, P. Helicase Subunit Cdc45 Targets the Checkpoint Kinase Rad53 to Both Replication Initiation and Elongation Complexes after Fork Stalling. Mol Cell 73, 562–573 e3 (2019).

71. Langston, L.D. et al. Mcm10 promotes rapid isomerization of CMG-DNA for replisome bypass of lagging strand DNA blocks. Elife 6(2017).

72. Looke, M., Maloney, M.F. & Bell, S.P. Mcm10 regulates DNA replication elongation by stimulating the CMG replicative helicase. Genes Dev 31, 291–305 (2017).

73. Mayle, R. et al. Mcm10 has potent strand-annealing activity and limits translocase-mediated fork regression. Proc Natl Acad Sci U S A 116, 798–803 (2019).

74. Ilves, I., Tamberg, N. & Botchan, M.R. Checkpoint kinase 2 (Chk2) inhibits the activity of the Cdc45/MCM2-7/GINS (CMG) replicative helicase complex. Proc Natl Acad Sci U S A 109, 13163–70 (2012).

75. Randell, J.C. et al. Mec1 is one of multiple kinases that prime the Mcm2-7 helicase for phosphorylation by Cdc7. Mol Cell 40, 353–63 (2010).

76. Sheu, Y.J., Kinney, J.B., Lengronne, A., Pasero, P. & Stillman, B. Domain within the helicase subunit Mcm4 integrates multiple kinase signals to control DNA replication initiation and fork progression. Proc Natl Acad Sci U S A 111, E1899–908 (2014).

77. Gilbert, C.S., Green, C.M. & Lowndes, N.F. Budding yeast Rad9 is an ATP-dependent Rad53 activating machine. Mol Cell 8, 129–36 (2001).

78. Donovan, S., Harwood, J., Drury, L.S. & Diffley, J.F. Cdc6p-dependent loading of Mcm proteins onto pre-replicative chromatin in budding yeast. Proc Natl Acad Sci U S A 94, 5611–6 (1997).

79. Liang, C. & Stillman, B. Persistent initiation of DNA replication and chromatin-bound MCM proteins during the cell cycle in cdc6 mutants. Genes Dev 11, 3375–86 (1997).

80. Ho, B., Baryshnikova, A. & Brown, G.W. Unification of Protein Abundance Datasets Yields a Quantitative Saccharomyces cerevisiae Proteome. Cell Syst 6, 192–205 e3 (2018).

